# Inferring disease architecture and predictive ability with LDpred2-auto

**DOI:** 10.1101/2022.10.10.511629

**Authors:** Florian Privé, Clara Albiñana, Julyan Arbel, Bogdan Pasaniuc, Bjarni J. Vilhjálmsson

## Abstract

LDpred2 is a widely used Bayesian method for building polygenic scores (PGS). LDpred2-auto can infer the two parameters from the LDpred model, the SNP heritability *h*^2^ and polygenicity *p*, so that it does not require an additional validation dataset to choose best-performing parameters. The main aim of this paper is to properly validate the use of LDpred2-auto for inferring multiple genetic parameters. Here, we present a new version of LDpred2-auto that adds an optional third parameter *α* to its model, for modeling negative selection. We then validate the inference of these three parameters (or two, when using the previous model). We also show that LDpred2-auto provides per-variant probabilities of being causal that are well calibrated, and can therefore be used for fine-mapping purposes. We also derive a new formula to infer the out-of-sample predictive performance *r*^2^ of the resulting PGS directly from the Gibbs sampler of LDpred2-auto. Finally, we extend the set of HapMap3 variants recommended to use with LDpred2 with 37% more variants to improve the coverage of this set, and show that this new set of variants captures 12% more heritability and provides 6% more predictive performance, on average, in UK Biobank analyses.

## Introduction

Most traits and diseases in humans are heritable. What differs is the genetic architecture of each trait that can be parameterized by three key terms: the heritability (i.e. the proportion of phenotypic variation explained by genetics), the polygenicity (i.e. the fraction of genomic variants that have a non-zero effect on the trait), and the causal effect distribution (i.e. how the effect size distribution varies across causal variants). Some phenotypes, such as schizophrenia or height, are highly heritable and highly polygenic (Sullivan *et al*., 2003; Yang *et al*., 2010; O’Connor *et al*., 2019; Trubetskoy *et al*., 2022). Causal effects are larger when a trait is more heritable, and smaller when it is more polygenic. The distribution of causal effects relative to their allele frequencies is often investigated through a single parameter, usually called *α* or *S*, to model the effect of negative selection on complex traits whereby variants with lower frequencies are expected to have higher causal effect sizes (Speed *et al*., 2012). Many methods have been developed to estimate the SNP heritability (referred to as *h*^2^ for brevity) and polygenicity (*p*), either globally for the whole genome or locally for specific regions of the genome, as well as *α*. These methods (non-exhaustively) include GCTA (*h*^2^, Yang *et al*. (2011)), BOLT-REML (*h*^2^ and *p*, Loh *et al*. (2015)), LD Score regression (*h*^2^, Bulik-Sullivan *et al*. (2015b)), FINEMAP (per-variant *p*, also called posterior inclusion probabilities PIPs, used for fine-mapping, Benner *et al*. (2016)), HESS (local *h*^2^, Shi *et al*. (2016)), LDAK-SumHer (*h*^2^, Speed and Balding (2019)), S-LD4M (*p*, O’Connor *et al*. (2019)), GRM-MAF-LD (*α*, Schoech *et al*. (2019)), SuSiE (PIPs, Wang *et al*. (2020)), SBayesS (*h*^2^, *p*, and a third parameter *S*, similar to *α*, Zeng *et al*. (2021)), and BEAVR (local *p*, Johnson *et al*. (2021)).

As previously shown by Daetwyler *et al*. (2008), *h*^2^ and *p* can also be used to determine how well we can predict a phenotype from using genetic variants alone 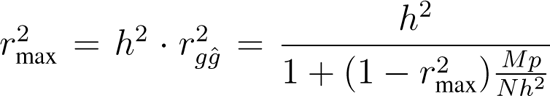, with where *r*^2^_max_ and *r*^2^_gg_ are the maximum achievable squared correlations between the genetic predictor and respectively the phenotype and its genetic component, *N* is the sample size, *M* is the number of variants, so *Mp* is the number of causal variants. Such genetic predictors are called polygenic scores (PGS), and are getting closer to being included as part of existing clinical risk models for diseases (Torkamani *et al*., 2018; Lambert *et al*., 2019; Kumuthini *et al*., 2022). LDpred2 is a widely used polygenic score method that can directly build PGS using resulting summary statistics from genome-wide associations studies (GWAS), making it highly applicable (Privé *et al*., 2020b; Pain *et al*., 2021; Kulm *et al*., 2021). LDpred2 is a Bayesian approach that uses the SNP heritability *h*^2^ and polygenicity *p* as parameters of its model. LDpred2-auto, one variant of LDpred2, can directly estimate these two parameters from the data, making it applicable even when no validation data is available for tuning these two model hyper-parameters (Privé *et al*., 2020b).

The main aim of this paper is to properly validate the use of LDpred2-auto for inferring multiple genetic parameters. Here we extend LDpred2-auto and show it is a reliable method for estimating *h*^2^ (global and local), *p* (also per-variant probabilities PIP used for fine-mapping purposes), and *α* (by extending its model to also include this third parameter). So, on top of providing competitive PGS, LDpred2-auto can also provide all these estimates of genetic architecture. Moreover, we show how it can now also reliably estimate the predictive ability *r*^2^ of PGS it derives, allowing for directly assessing the usefulness of the derived PGS, without requiring an independent test set. An overview of what LDpred2-auto can provide is presented in Figure 1. Finally, we extend the set of HapMap3 variants recommended to use with LDpred2, which enables us to capture around 12% more SNP heritability and achieve around 6% more predictive performance *r*^2^, on average, in UK Biobank analyses. We call ‘HapMap3+’ this extended set of 1,444,196 SNPs, and recommend to use it when power is sufficient.

**Figure 1:**
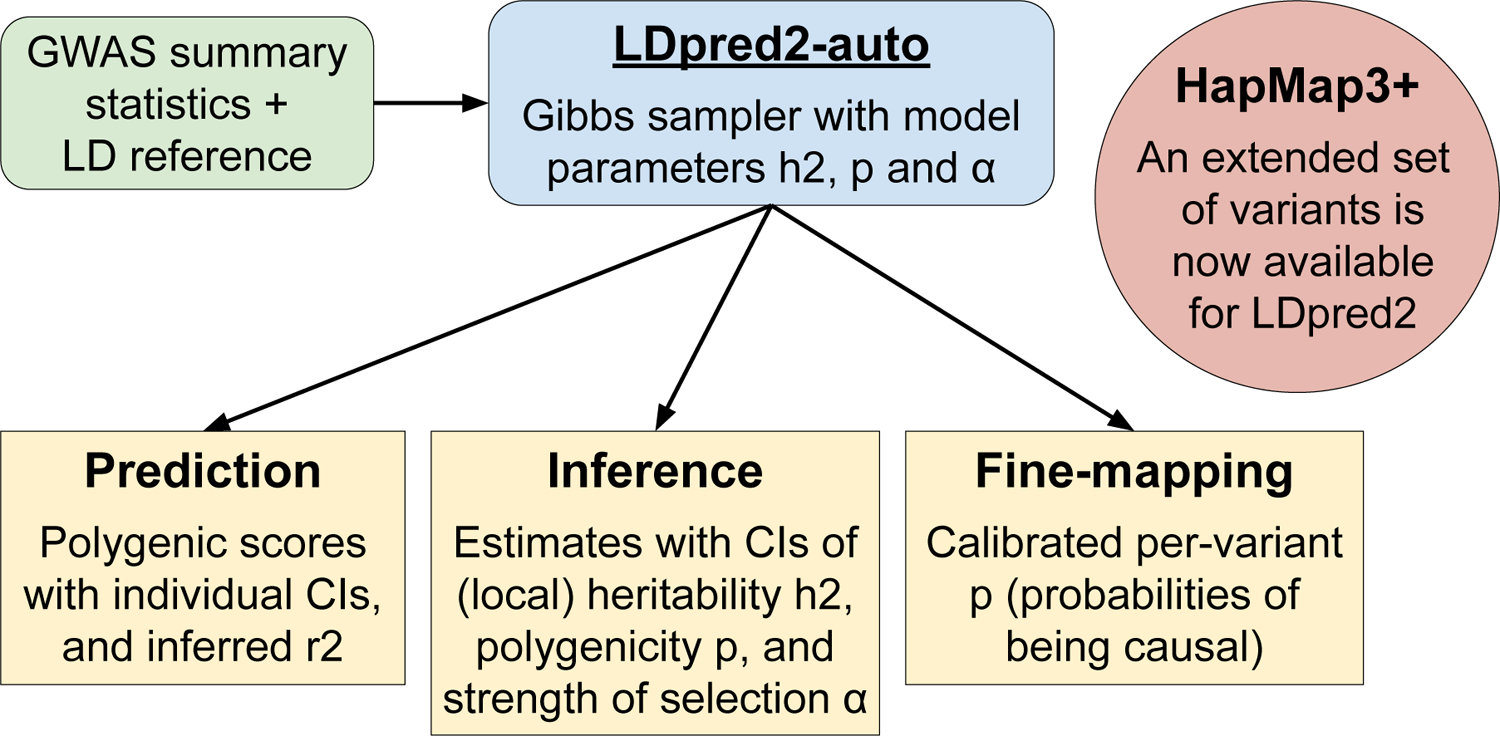
Overview of what LDpred2-auto can now provide. For individual CIs of polygenic scores, please refer to the work of Ding *et al*. (2022, 2023). HapMap3+ is the new extended set of 1,444,196 SNPs we introduce here (Methods), and now recommend to use for LDpred2 when power is sufficient. CI means “confidence interval” (often called “credible interval” in a Bayesian setting).

## Results

### Validating the inference with simulations

For simulations, we use the UK Biobank imputed data (Bycroft *et al*., 2018). We use 356,409 individuals of Northwestern European ancestry and 322,805 SNPs across seven chromosomes (Methods). We first simulate continuous phenotypes using function `snp_simuPheno` from R package bigsnpr (Privé *et al*., 2018), varying three parameters: the SNP heritability *h*^2^, the polygenicity *p* (i.e. the proportion of causal variants), and the parameter *α* in equation (2) that controls the relationship between minor allele frequencies and expected effect sizes. This function first picks a proportion *p* of causal variants at random, samples effect sizes *γ* using the variance component parameterized by *α* and then scales the effect sizes so that the genetic component *Gγ* has a variance *h*^2^, where *G* is the genotype matrix. Finally, some Gaussian noise is added so that the final phenotype has a variance of 1. Then, we run a GWAS to obtain summary statistics using *N* individuals (either the 200,000 dedicated to this, or a random subset of 20,000), using fast linear regressions implemented in `big_univLinReg` from R package bigstatsr (Privé *et al*., 2018). Finally, we run LDpred2-auto with and without the option `allow_jump_sign`, which was proposed in Privé *et al*. (2022a) for robustness (when disabled, it prevents effect sizes from changing sign without going through 0 first), and with and without the new model including a third parameter *α* (new option `use_MLE`, Methods). LDpred2-auto is run with 50 Gibbs sampler chains with different starting values for *p* (from 0.0001 to 0.2, equally spaced on a log scale). Then some of these chains are filtered out for quality control (Methods).

First, LDpred2-auto generally reliably infers the three parameters from its model, i.e. the SNP heritability *h*^2^, polygenicity *p*, and *α* (Figures 2 and S1–S5). Compared to LD Score regression, heritability estimates are as precise when power is low, and much more precise when power is large, especially for small polygenicity values (Figures 2 and S1). When power is low (e.g. *N* = 20000 and *h*^2^ = 0.01), LD-pred2_noMLE (with only two model parameters, as in previous versions of LDpred2-auto) and SBayesS are both over-confident in their estimate of the heritability (i.e. CIs are small, and do not contain the true parameter value), and LDpred2_jump overestimates the heritability (Figure 2). LDpred2_nojump looks very reliable for estimating *h*^2^. When power is low, LDpred2-auto can overestimate the polygenicity when the true value is very small (e.g. *p* = 0.0005), and underestimate it when the polygenicity is large (e.g. *p* = 0.1, Figure S2). SBayesS, which uses a similar model with the same three parameters, often overestimates the polygenicity, especially when *p ≤* 0.02. The *α* estimate of LDpred2-auto (with the new 3-parameter model) can become very unprecise when power is too low, which can be detected by a small number of chains kept from LDpred2-auto, while estimates from SBayesS are often over-confident with small CIs that do not include the true simulated values (Figure S4). Finally, we also investigate new ways of post-filtering chains in LDpred2-auto, for both prediction and inference (“LDpred2_newfilter”, Methods); results remain practically unchanged (Figures 2 and S1–S5).

**Figure 2:**
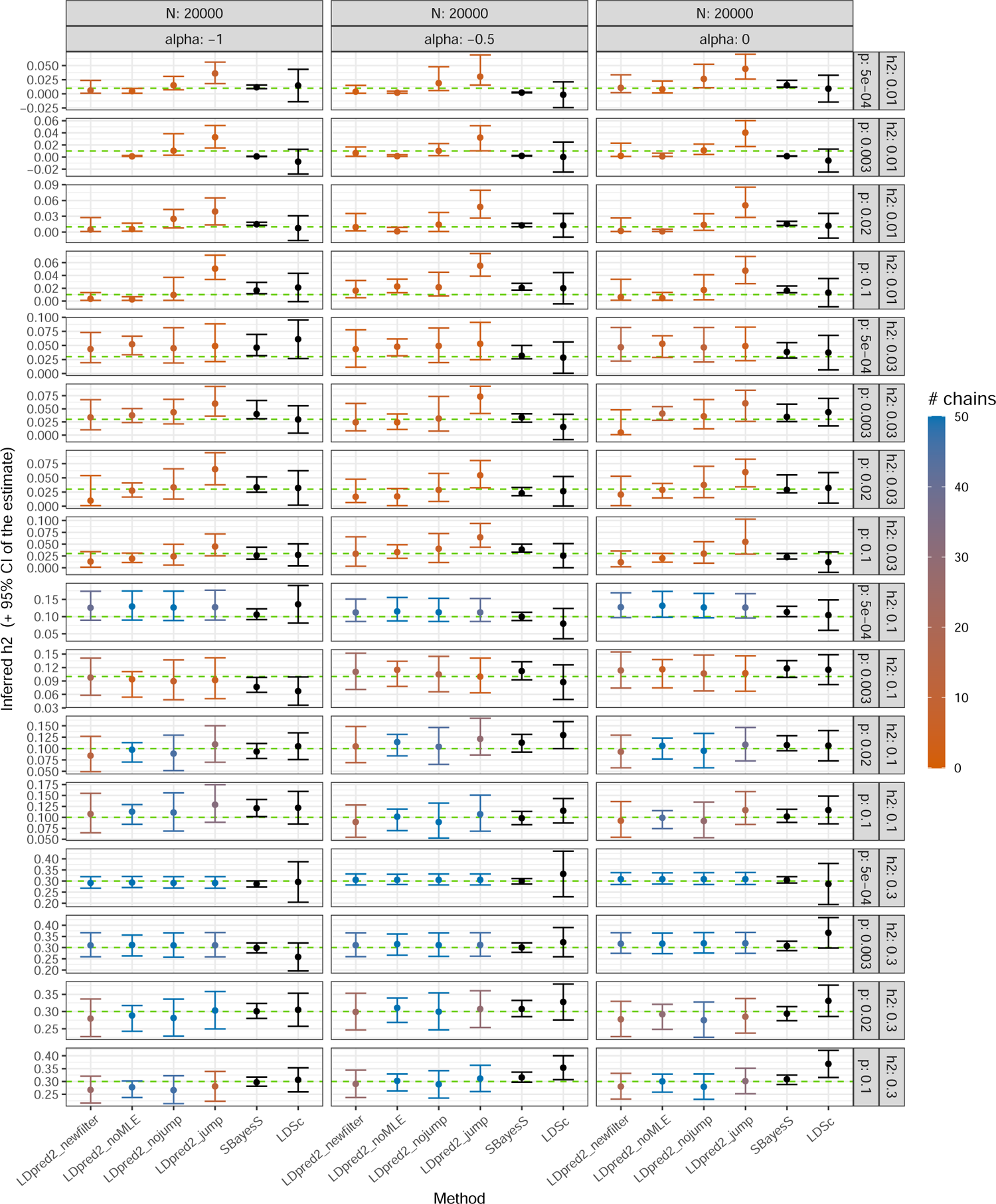
Inferred SNP heritability *h*^2^ in simulations with continuous outcomes and N=20K. Horizontal dashed lines represent the true simulated values. For LDpred2-auto, suffix “nojump”/“jump” refers to using `allow_jump_sign = FALSE/TRUE` (and `use_MLE = TRUE`), and “noMLE” refers to using `use_MLE = FALSE` (and `allow_jump_sign = FALSE`), and “newfilter” is similar to “no-jump” but uses a different post-filtering of chains (Methods). Note that the recommended option is to use `allow_jump_sign = FALSE` (Privé *et al*., 2022a). The 95% confidence intervals for the LDpred2-auto and SBayesS estimates are obtained from the 2.5% and 97.5% quantiles of all the *h*^2^ estimates from the iterations (after burn-in) of the chains kept (note that only one chain is used and kept in SBayesS). The 95% confidence interval for the LD Score (LDSc) regression estimate is obtained as *±*1.96 of its standard error. Colors for LDpred2-auto models represent the number of chains kept (out of 50).

Then, LDpred2-auto can also infer per-variant probabilities of being causal and local per-block heritability estimates, which are well calibrated (Figures S6 and S7). We recall that calibrated per-variant probabilities of being causal (also known as posterior inclusion probabilities, PIPs) can be used for fine-mapping purposes (Wang *et al*., 2020). LDpred2-auto provides PIPs that are more calibrated than with e.g. SuSiE-RSS (Zou *et al*., 2022), which we run assuming there are 10 causal variants per LD block by default (Figure S8). Finally, LDpred2-auto can also be used to reliably infer the predictive performance *r*^2^ of its resulting polygenic score, directly from within the Gibbs sampler (Methods), even when power is low, and we show it works with results from SBayesS’s Gibbs sampler as well (Figures S9 and S10).

We then run simulations with binary outcomes where the simulated continuous liabilities are transformed to binary outcomes using a threshold corresponding to the prevalence. Results are very similar as with the continuous phenotypes above (Figures S11–S14), and are similar whether we use either a linear regression GWAS and the total sample size *N*, or a logistic regression GWAS and the effective sample size (i.e. *N*_eff_ = 4/(1/*N*_case_ + 1/*N*_control_)). The main difference is that the *h*^2^ and *r*^2^ estimates must be transformed to the liability scale (Lee *et al*., 2011), where *K*_GWAS_ = 0.5 should be used for transforming estimates of *h*^2^ and *r*^2^ when using *N*_eff_ in inference methods (Grotzinger *et al*., 2023).

### Genetic architectures of 248 phenotypes from the UK Biobank

We use the same 356,409 unrelated individuals of Northwestern European ancestry as in the simulations. To form the test set, we randomly select 50,000 of these, while the other 306,409 are used to run a GWAS using linear regression (with function `big_univLinReg` from R package bigstatsr) for each of all 248 phenotypes and using eight covariates (Methods). We first use the set of 1,054,330 HapMap3 variants recommended to use for LDpred2 (Privé *et al*., 2020b). Here, if not otherwise specified, we use options `use_MLE = TRUE` (i.e. the new 3-parameter model and sampling scheme) and `allow_jump_sign = FALSE`.

Consistent with simulations, inferred SNP heritability *h*^2^ estimates from LDpred2-auto closely match with those from LD Score regression, while generally being more precise, especially for phenotypes with a small polygenicity (Figure S15). Note that these *h*^2^ estimates (and later the *r*^2^ estimates) have not been transformed to the liability scale (i.e. are on the observed scale). Most phenotypes have an estimated polygenicity *p* between 0.001 and 0.04; these have therefore a very polygenic architecture, but not an infinitesimal one (Figure S16). Most phenotypes have an estimated *α* between −1.0 and −0.4 with a mode at −0.65 (Figure S17). As for the inferred predictive performance *r*^2^, they are highly consistent with the ones derived from the test set; only for standing height are they overestimated (Figure 3). Heritability estimates for height are probably overestimated as well since we use similar formulas for estimating *h*^2^ and *r*^2^ (Methods), and because the SNP heritability estimate *h*^2^ for standing height is higher than values reported in the literature (also see Section “Application to height”).

**Figure 3:**
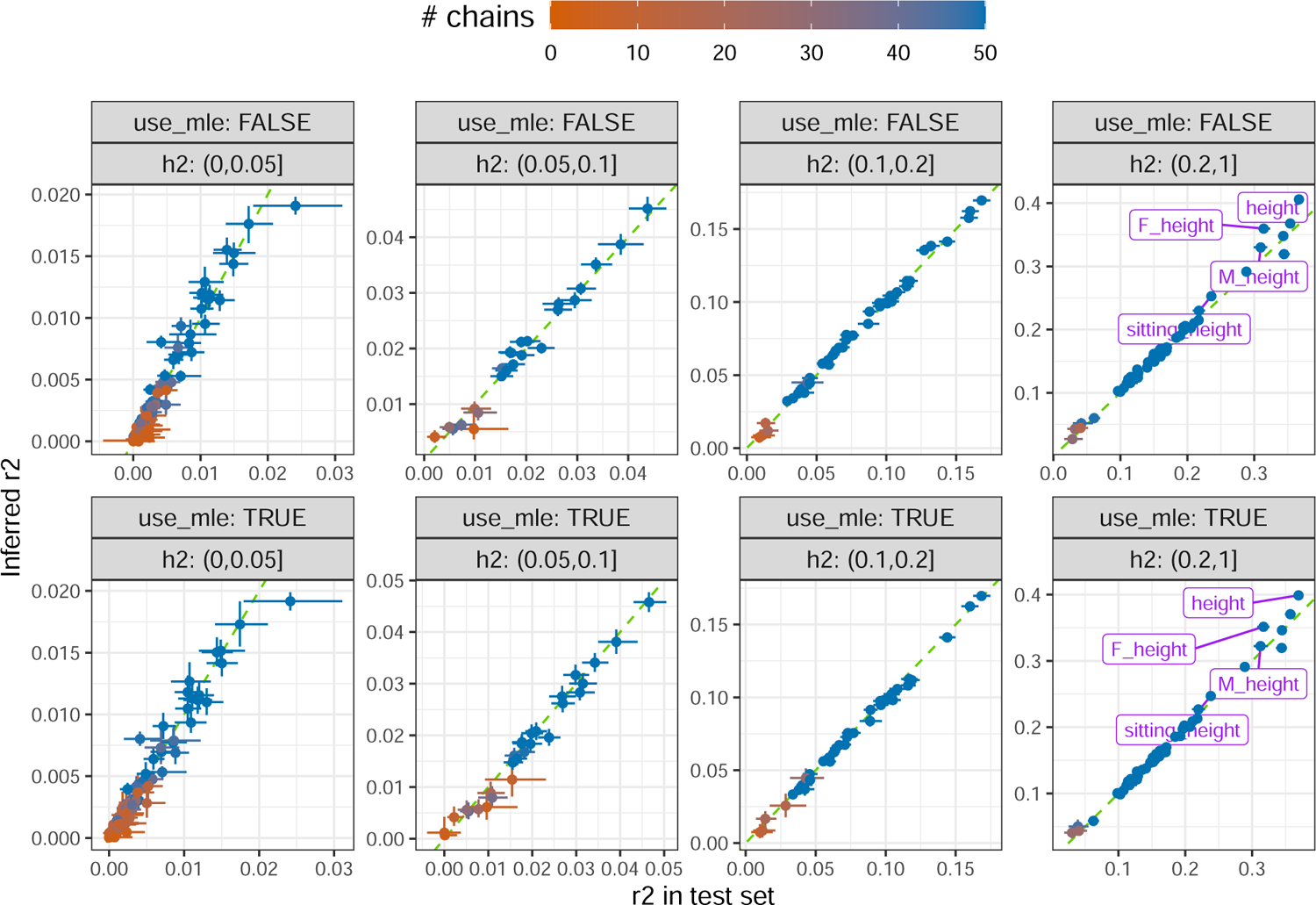
Inferred predictive performance *r*^2^ from the Gibbs sampler of LDpred2-auto versus the ones obtained in the test set, for all 248 phenotypes defined from the UK Biobank. These are stratified by the polygenicity estimated from LDpred2-auto. Green dashed lines represent the 1:1 line. The 95% confidence interval for the LDpred2-auto estimate is obtained from the 2.5% and 97.5% quantiles of all the *r*^2^ estimates from the iterations (after burn-in) of the chains kept. The 95% confidence interval for *r*^2^ in the test set is obtained from bootstrap. Colors represent the number of chains kept (out of 50). “F_height” and “M_height” use females and males only, respectively (in both GWAS and test sets).

To investigate whether estimates from LDpred2-auto are robust to some misspecifications, we test using two alternative LD references (Methods). Using a smaller number of individuals for computing the LD matrix results in a slightly overestimated *p* and *h*^2^ (and *r*^2^), while the *α* estimate remains consistent, and the predictive performance in the test set remains mostly similar, except for three phenotypes for which none of the LDpred2-auto chains is usable (Figure S18). When using an LD reference from an alternative population (South Europe instead of North-West Europe), *p*, *h*^2^, and *r*^2^ are slightly overestimated as well, and a few phenotypes have lower predictive performance while there are four phenotypes for which none of the LDpred2-auto chains is usable (distinct from the previous three, Figure S19). Interestingly, using the previous approach (`use_MLE = FALSE`) seems to provide more robust results, where we can always get some chains not to diverge (and therefore get non-zero predictive performance) for the seven (three and four) phenotypes mentioned before when using the previous alternative LD references (Figure S20).

Then, we investigate the effect of disabling the LDpred2-auto parameter `allow_jump_sign` (for extra robustness as shown in Privé *et al*. (2022a)) on the estimates from LDpred2-auto. Consistent with simulations, *p* estimates from LDpred2-auto are conservatively lower than when allowing effects to “jump” sign (i.e. normal sampling, Figure S21). *h*^2^ estimates can also be slightly lower, while *α* estimates are broadly consistent. As for predictive performance *r*^2^ (on the test set), they are similar, suggesting there is no problem of robustness here (when using the normal LD reference) and normal sampling can be used in this case (Figure S21).

Finally, we investigate different transformations to apply to some continuous phenotypes used here. Indeed, 49 of the phenotypes used here seem log-normally distributed or heavy-tailed (when visualizing their histogram); we therefore log-transform them. However, we do investigate alternative transformations here to decide which one should be preferred and to check how this impacts the inference and prediction from LDpred2-auto. We first compare to using raw (untransformed) phenotypes in Figure S22; estimates of *p* and *α* are highly consistent. However, *h*^2^ estimates and predictive performance *r*^2^ (in the test set) are generally larger with the log-transformation; it probably makes sense to transform these phenotypes. We then compare to using the rank-based inverse normal (RIN) transformation in Figure S23; estimates for *p* and *α* are also highly consistent. Except for bilirubin and lipoprotein(a) concentration, generally higher *h*^2^ estimates and predictive performance *r*^2^ are obtained with the RIN-transformation than the log-transformation.

### More heritability and predictive accuracy with new set of variants

Here we use the same individuals as in the previous section. We investigate using the extended set of HapMap3 variants proposed here, HapMap3+ (Methods), which includes ∼37% more variants on top of HapMap3 variants recommended to use for LDpred2 (i.e. 1,054,330 + 389,866 variants), to improve the genome coverage of this set. As expected, compared to HapMap3, higher *h*^2^ (average increase of 12.3% [95% CI: 10.8, 13.7]) and lower *p* (decrease of 11.5% [10.7, 12.3]) estimates are obtained with this extended set HapMap3+ (Figure 4). This is consistent with higher predictive performance *r*^2^ in the test set (increase of 6.1% [4.1, 8.2]). In particular, a much larger *h*^2^ estimate is obtained for lipoprotein(a) concentration (0.508 [0.502, 0.514] instead of 0.324 [0.320, 0.329]), which is also reflected in a larger predictive performance (*r*^2^ in the test set of 0.516 [0.508, 0.524] instead of 0.344 [0.335, 0.353]). Interestingly, when using this extended set of HapMap3 variants, more chains are kept on average, which is a sign of better convergence of the models (Figure S24). However, running LDpred2 with this extended set of variants takes around 50% more time; yet, we remind the reader that LDpred2 has been made much faster in Privé *et al*. (2022a), and now runs in less than one hour for 50 chains parallelized over 13 cores (Figure S25), instead of 4–12 hours before.

**Figure 4:**
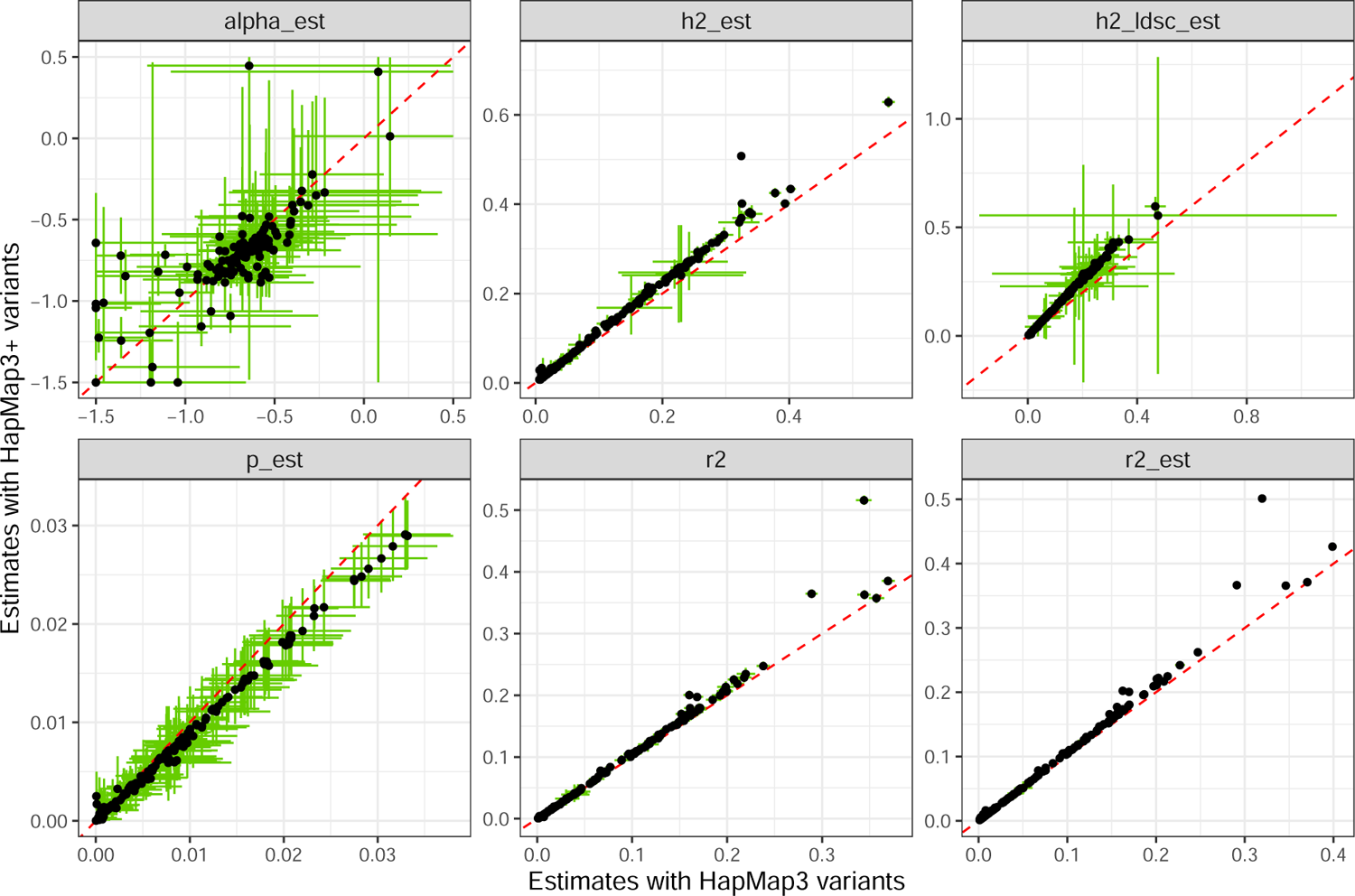
LDpred2-auto estimates for UKBB phenotypes using either the HapMap3 or HapMap3+ sets of variants. Only 154 phenotypes with more than 25 chains kept when using the HapMap3 variants are represented here. Red dashed lines represent the 1:1 line. The 95% confidence interval for the LDpred2-auto estimate (in green) is obtained from the 2.5% and 97.5% quantiles of all the estimates from the iterations (after burn-in) of the chains kept. The 95% confidence interval for *r*^2^ in the test set is obtained from bootstrap.

### Local heritability and polygenicity

In this section, we use the extended set of variants constructed here, HapMap3+, for which we define 431 independent LD blocks (Methods). We compute local per-block *h*^2^ estimates, and report the UKBB phenotypes for which one block contributes to at least 10% of the total heritability of all blocks in Figure S26. For lipoprotein(a) concentration, “red hair” and “disorders of iron metabolism” (phecode 275.1), almost all heritability comes from one LD block only. We also perform the same analysis with external GWAS summary statistics for 90 cardiovascular proteins (Folkersen *et al*., 2020); 22 (resp. 8) of them have at least 50% (resp. 80%) of their heritability explained by a single block (Figure S27).

Across 169 UKBB phenotypes with more than 25 chains kept, we compute the median heritability per block, and compare it to the number of variants in these blocks; the median heritability explained by a block of variants is largely proportional to the number of variants in this block (Figure S28). The outlier block explaining a much larger heritability contains the HLA region. Across the same phenotypes, we then compute per-variant median probabilities of being causal, and report them in a Manhattan-like plot in Figure S29. Some variants in multiple small regions across the genome have a larger probability of being causal across many phenotypes; interestingly, these are mapped to genes that are known to be associated with many different traits (up to more than 300) in the GWAS Catalog (Buniello *et al*., 2019). To verify that this is not driven by population structure, we compute pcadapt chi-square statistics that quantify whether a variant is associated with population structure (Privé *et al*., 2020c); the log-statistics have a small negative correlation of −5.5% with the probabilities of being causal. To verify that this does not correspond to regions of low LD, we compute LD scores; the median probabilities of being causal have a small correlation of 11.6% with the LD scores and of −6.8% with the log of LD scores.

### Application to height

Here we run three LDpred2-auto models for height, one from the same 305K training UKBB individuals used before (with available height, out of 306K), one based on 100K UKBB individuals (as a random subset of the previous 305K), and one from a large GWAS meta-analysis of 1.6M individuals of European genetic ancestries (Yengo *et al*., 2022). We first infer the genetic ancestry proportions of individuals included in the meta-analysis using the method proposed in Privé (2022), and find that 81.9% are from N.W. Europe, 9.5% from E. Europe, 6.5% from Finland, 1.5% of Ashkenazi genetic ancestry, 0.3% from S.W. Europe, and 0.2% from W. Africa. For this set of external GWAS summary statistics, we therefore use the same N.W. European LD matrix as used in UKBB analyses. Note that we use the HapMap3+ set of 1,444,196 SNPs here, however, for the GWAS meta-analysis, only 1,013,499 SNPs (out of 1,373,020) are overlapping with HapMap3+ and passing quality control.

As expected (Loh *et al*., 2018), intercepts from LD Score regression are increasing with sample size: 1.02 (SE: 0.008) with N=100K, 1.11 (0.015) with N=305K, and 2.31 (0.068) with N=1.6M. SNP heritability estimates are 64.6% (SE: 2.7), 59.7% (2.2), and 39.2% (1.7) with LD Score regression, respectively, and 60.2% [95% CI: 57.2, 63.2], 63.2% [62.0, 64.4], and 54.2% [53.9, 54.5] with LDpred2-auto.

As expected, predictive performance *r*^2^ (estimated from the Gibbs sampler) are increasing with sample size, with 29.6% [28.7, 30.5], 42.7% [42.2, 43.1], and 47.0% [46.8, 47.1], respectively. Note that these *r*^2^ estimates are probably overestimated by the same margin as the (SNP heritability) *h*^2^ estimates, and correspond to ∼49%, ∼67.5%, and ∼87% of *h*^2^, respectively. Even though there are 1.6M individuals in the meta-analysis, the predictive performance corresponds to around 87% of the SNP heritability only, therefore an even larger sample size is required to be able to better predict height. Polygenicity estimates from LDpred2-auto are increasing with sample size with 1.2% [1.0, 1.5], 2.3% [2.0, 2.5], and 5.9% [5.6, 6.3], consistent with results of simulations with a large polygenicity (p=10%). Therefore, we estimate that height has at least 50,000 causal variants. These results are similar irrespective of whether `allow_jump_sign` is used or not, which is surprising to us. We also identify 1753 SNPs with a probability of being causal larger than 95% (fine-mapping), which are spread over the entire genome (Figure S30). As for *α* estimates from LDpred2-auto, they remain consistent, with −0.71 [−0.75, −0.67], −0.74 [−0.76, −0.72], and −0.78 [−0.82, −0.76], respectively. Finally, we compute per-annotation heritability estimates to investigate functional enrichment. We perform this analysis using 50 non-flanking binary annotations from the baselineLD v2.2 model (Finucane *et al*., 2018). Heritability enrichments are rather modest, ranging from 0.7 to 2.5 with a GWAS sample size of N=305K, and of slightly smaller magnitude with N=100K and slightly larger magnitude with N=1.6M (Figure S31).

### Application to other external GWAS summary statistics

A description of the eight external GWAS summary statistics used is provided in Table 1; these do not include UKBB individuals. Quality control of these summary statistics is performed as described in Privé *et al*. (2022a). We run LDpred2-auto using either the HapMap3 or HapMap3+ variants, with either the new or previous sampling and model (via parameter `use_MLE`). Because of the increased mismatch between external GWAS summary statistics and the LD reference we use here (compared to in simulations and UKBB analyses), we also explore multiple values for parameter `coef_shrink`, which is a multiplicative coefficient for shrinking/regularizing off-diagonal elements of the LD matrix in LDpred2-auto. Note that, to transform *r*^2^ and *h*^2^ estimates to the liability scale (except for vitamin D, which is a continuous trait), we use the prevalence in the UK Biobank as the population prevalence, which may be slightly biased Fry *et al*. (2017); van Alten *et al*. (2022). Results are presented in Figure S32. In terms of predictive performance, using the HapMap3+ variants provides equal or better predictive performance compared to using the HapMap3 variants, except for type 1 diabetes (T1D); therefore it seems more useful to use this new set of variants for larger sample sizes. Using the new sampling scheme and model (with option `use_MLE`) provides equal or better predictive performance except for vitaminD, but proves to be less robust, especially when using `coef_shrink` close to 1 (low or no regularization of the LD matrix).

**Table 1:** Summary of external GWAS summary statistics used. These do not include UKBB individuals. Note that some of them may contain a substantial amount of non-European genetic ancestry (e.g. >20%) for CAD.

| Trait | GWAS citation | Effective GWAS sample size | # GWAS variants | # matched variants with HapMap3+ |
| --- | --- | --- | --- | --- |
| Asthma | Demenais *et al.* (2018) | 67,341.2 | 2,001,280 | 946,092 |
| Breast cancer (BRCA) | Michailidou *et al.* (2017) | 254,862.6 | 11,792,542 | 1,411,710 |
| Coronary artery disease (CAD) | Nikpay *et al.* (2015) | 129,014.3 | 9,455,778 | 1,325,052 |
| Depression (MDD) [without UKBB] | Wray *et al.* (2018) | 168,040.2 | 9,874,289 | 1,314,499 |
| Prostate cancer (PRCA) | Schumacher *et al.* (2018) | 135,316.1 | 20,370,946 | 1,411,710 |
| Type 1 diabetes (T1D) | Censin *et al.* (2017) | 13,497.6 | 8,996,866 | 1,127,489 |
| Type 2 diabetes (T2D) | Scott *et al.* (2017) | 72,143.0 | 12,056,346 | 1,408,283 |
| Vitamin D | Jiang *et al.* (2018) | 79,366 | 2,579,296 | 1,028,171 |

Maximum *r*^2^ are achieved at different LD regularization coefficients `coef_shrink` across phenotypes, reflecting possible substantial mismatches between the GWAS summary statistics and LD reference used. However, results are virtually unchanged when regularizing the LD matrix (“hm3_plus_regul”) by multiplying correlations between variants *i* and *j* by exp(*−*0.5 *· d_i,j_*), where *d_i,j_* is the distance in cM between the two variants (similarly to as proposed in Wen and Stephens (2010)), which is surprising to us. As for estimates of *h*^2^ and *r*^2^ (inferred from the Gibbs sampler), they tend to increase with smaller values of `coef_shrink`. This is also the case for estimates of *p* when using `use_MLE = TRUE`. Estimates of *α* are largely constant but can become very small (capped at −1.5) in the case of robustness issues when using e.g. `use_MLE = TRUE` and almost no regularization on the LD matrix (i.e. `coef_shrink` close to 1).

## Discussion

LDpred2-auto was originally developed for building polygenic scores (Privé *et al*., 2020b). Here we have extended the LDpred2-auto model and shown that it can be used to reliably infer genetic architecture parameters such as the SNP heritability (both genome-wide and more locally), polygenicity (and per-variant probabilities of being causal, also known as posterior inclusion probabilities PIPs, useful for fine-mapping), and the selection-related parameter *α*. We remind readers that LDpred2 can also be used to infer the uncertainty of individual polygenic scores (Ding *et al*., 2022). We also present a new way to infer the out-of-sample predictive performance *r*^2^ of the resulting PGS, assuming the target sample has the same genetic ancestry as the GWAS used for training. Others have proposed to estimate *r*^2^ from a reference genotype set (Mak *et al*., 2017) or from an additional set of external GWAS summary statistics and LD reference (Pattee and Pan, 2020; Witteveen *et al*., 2022); here we only use the summary statistics that we input to LDpred2-auto. Results across 248 phenotypes demonstrate that most of these phenotypes are very polygenic, yet do not have an infinitesimal architecture (i.e. not all variants are causal); this is consistent with LDpred2-inf (assuming an infinitesimal architecture, i.e. *p* = 1) generally providing lower predictive performance than LDpred2-grid (testing different values for parameter *p*) or LDpred2-auto (estimating *p*, Privé *et al*. (2020b)). We also obtain widespread signatures of negative selection with most *α* estimates between −1.0 and −0.4, consistent with previous findings (Zeng *et al*., 2021). However, when looking at the heritability enrichment of several functional annotations for height, we obtain much smaller magnitudes than stratified LD Score regression (S-LDSC, Finucane *et al*. (2018)). For example, Yengo *et al*. (2022) report fold enrichments of more than 10x for e.g. coding and conserved variants, while we get less than 2x. This is partly due to LDpred2-auto estimates being more conservative as they are shrunk towards no enrichment (the prior), however we do use a very large sample size here so that the prior should not matter much. Another possible reason comes from using variants that capture the causal effects by LD, while these tagging variants may be annotated differently from the causal variants, which could cause functional enrichments to be diluted (Zheng *et al*., 2022). We also note that this heritability partitioning is performed after running LDpred2-auto for each annotation independently, therefore, unlike S-LDSC, the LDpred2-auto heritability partitioning does not depend on the set of annotations used.

Here we have also extended the set of HapMap3 variants recommended to use with LDpred2, making it 37% larger to offer a better coverage of the genome. Since we used individuals of diverse ancestries for computing the pairwise variant correlations used for constructing this extended set of variants, we expect this new set of variants to be beneficial across diverse ancestries. Increasing the genome coverage enables us to capture more of the heritability of phenotypes and therefore reduce the missing heritability, i.e. the difference between the family-based heritability and the SNP-based heritability. Using the new HapMap3+ set also improves predictive performance by an average of 6.1% in UKBB analyses here, and particularly for lipoprotein(a) concentration with an *r*^2^ of 0.516 instead of 0.344. However, we note that we are able to achieve an *r*^2^ of 0.677 [0.671, 0.682] when using the penalized regression implementation of Privé *et al*. (2019) on the UKBB individual-level data while restricting to all variants within a 1Mb window of the *LPA* gene. This means that this extended SNP set is still not tagging all variants perfectly, and that it might be preferable to use a more localized set of variants for phenotypes for which most of the heritability is contained in a single region of the genome. When using external GWAS summary statistics, using HapMap3+ instead of HapMap3 variants was more beneficial for larger sample sizes. Our intuition and conclusion is that using more variants is beneficial when power is sufficient; however, when power is low (e.g. small *N*, small *h*^2^, and/or large *p*), it may be actually be detrimental.

Our proposed method has limitations. First, when power is low (i.e. when *Nh*^2^*/p* is low), estimates of *α* and *p* become less reliable. Therefore, we do recommend to use the 3-parameter model (with *α*), but only when power is sufficient. However, estimates of *h*^2^ and *r*^2^ seem always reliable, except for height for which they are probably overestimated. We think this is likely due to assortative mating (Border *et al*., 2022; Yengo *et al*., 2022). Second, the *h*^2^ from LDpred2-auto is also slightly overestimated when using a small LD reference panel or when the reference panel does not closely match with the ancestry of the GWAS summary statistics. Future work could focus on correcting these issues. Third, when using external GWAS summary statistics, it is often beneficial to regularize the LD matrix (via parameter `coef_shrink`, especially when using the new sampling and model), however it leads to an increased estimation for e.g. *r*^2^ and *h*^2^. Future work could focus on correcting these estimates, and also on identifying the best shrinkage regularization coefficient to apply based on e.g. some distance metric (mismatch) between GWAS summary statistics and the LD reference used. Fourth, because we use a limited set of variants as input for LDpred2, causal variants identified by LDpred2-auto are probably tagging variants that are highly correlated with unobserved causal variants close by.

Nevertheless, LDpred2-auto users can now get much from running a single method. The reliable estimates provided by LDpred2-auto are very encouraging to further extend LDpred2-auto in multiple directions. As future research directions, we are interested in using LDpred2-auto for GWAS summary statistics imputation (Rüeger *et al*., 2018; Julienne *et al*., 2019), for genetic correlation estimation (Bulik-Sullivan *et al*., 2015a; Shi *et al*., 2017; Speed and Balding, 2019; Frei *et al*., 2019; Werme *et al*., 2022), multi-ancestry prediction and inference (Brown *et al*., 2016; Shi *et al*., 2020; Ruan *et al*., 2022; Lu *et al*., 2022), as well as extending it to use more variants and learn from functional annotations (Zhang *et al*., 2021; Márquez-Luna *et al*., 2021; Zheng *et al*., 2022).

## Materials and Methods

### Data for simulations

For simulations, we use the UK Biobank imputed (BGEN) data, read as allele dosages with function `snp_readBGEN` from R package bigsnpr (Bycroft *et al*., 2018; Privé *et al*., 2018). We use the set of 1,054,330 HapMap3 variants recommended to use for LDpred2 (Privé *et al*., 2020b). Since we run lots of different models, we restrict the simulations to chromosomes 3, 6, 9, 12, 15, 18 and 21, resulting in a set of 322,805 SNPs. We restrict individuals to the ones used for computing the principal components (PCs) in the UK Biobank (field 22020). These individuals are unrelated and have passed some quality control such as removing samples with a missing rate on autosomes larger than 0.02, having a mismatch between inferred sex and self-reported sex, and outliers based on heterozygosity (more details can be found in section S3 of Bycroft *et al*. (2018)). To get a set of genetically homogeneous individuals, we compute a robust Mahalanobis distance based on the first 16 PCs (field 22009) and further restrict individuals to those within a log-distance of 4.5 (Privé *et al*., 2020a). This results in 356,409 individuals of Northwestern European ancestry. We randomly sample 200,000 individuals to form a training set (to run the GWAS), and use the remaining individuals to form a test set (to evaluate the predictive models).

### Data for the UK Biobank analyses

We use the set of 1,054,330 HapMap3 variants recommended to use for LDpred2 (Privé *et al*., 2020b), and the same 356,409 individuals of Northwestern European ancestry as in the simulations. We randomly sample 50,000 individuals to form a test set (to evaluate the predictive models), and use the remaining individuals to form a training set (to run the GWAS).

We construct and use the same phenotypes as in Privé *et al*. (2022b). About half of these consists of phecodes mapped from ICD10 and ICD9 codes using R package PheWAS (Carroll *et al*., 2014; Wu *et al*., 2019). The other half consists of phenotypes defined in UKBB fields based on manual curation (Privé *et al*., 2022b). As covariates, we first recompute PCs for the homogeneous subset of individuals previously defined using function `snp_autoSVD` from R package bigsnpr and keep four PCs based on visual inspection (Privé *et al*., 2018, 2020a). We also use sex (field 22001), age (field 21022), birth date (combining fields 34 and 52) and deprivation index (field 189) as additional covariates (to a total of eight).

We use the LD matrix with independent LD blocks computed in Privé *et al*. (2022a). We design two other LD matrices: one using a smaller random subset of 2000 individuals from the previously selected ones (which we call “hm3_small”), and one based on 10,000 individuals from around South Europe by using the “Italy” center defined in Privé *et al*. (2022b) (“hm3_altpop”). We apply the optimal algorithm developed in Privé (2021) to obtain independent LD blocks, as recommended in Privé *et al*. (2022a). We finally define a fourth LD reference by extending the set of HapMap3 variants (see next Methods section) and using 20,000 individuals from the previously selected ones (“hm3_plus”).

### Extending the set of HapMap3 variants used

The HapMap3 variants generally provide a good coverage of the whole genome. We recall that the set of 1,054,330 HapMap3 variants recommended to use for LDpred2 (Privé *et al*., 2020b) is a subset of the original set of HapMap3 variants, which does not include duplicated positions (e.g. multi-allelic variants), nor ambiguous variants (e.g. ‘A’ and ‘T’, or ‘C’ and ‘G’), and which includes SNPs only (e.g. no indel). Here we propose to extend this set of 1,054,330 HapMap3 variants to make sure many genetic variants are well tagged by the extended set. To design this new set, we first read all variants from the UK Biobank (UKBB) with a minor allele frequency (MAF) larger than 0.005 in the whole data (i.e. the MAF from the MFI files). There are around 11.5M such variants. Then we restrict to unrelated UKBB individuals which are *not* listed as White British (field 22006) and use these individuals of diverse ancestries (Bycroft *et al*., 2018; Privé, 2022) to compute all pairwise correlations between variants within a 1 Mb distance, restricting to squared correlations larger than 0.3. Finally, we design an algorithm which aims at maximizing the tagging of all these variants read. We want to maximize where *j* spans the whole set of variants read, while *k* spans the variants kept in the new set, which we call HapMap3+. We start by including all previously used HapMap3 variants. Then, for the sake of simplicity, we use a greedy approach, where we repeatedly include the variant which increases this sum most, until no variant improves it by more than 2. Note that we only allow non-ambiguous SNPs to be included. This results in an extended set of 1,444,196 SNPs, of which we compute the LD matrix (within a 3 cM window) and apply the optimal algorithm developed in Privé (2021) to obtain 431 independent LD blocks. Since we use individuals of diverse ancestries for computing the pairwise variant correlations used for constructing this extended set of variants, we expect this new set of variants to be beneficial across diverse ancestries.

### New optional model and inference with LDpred2-auto

LDpred2 originally assumed the following model for effect sizes,

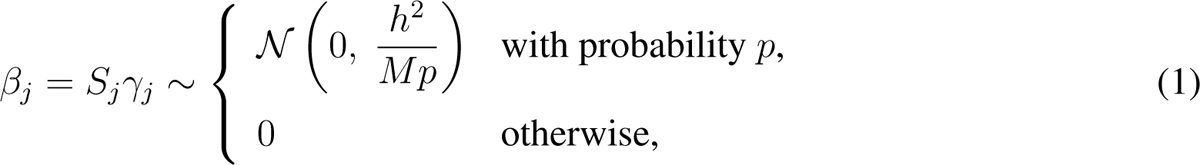

where *p* is the proportion of causal variants, *M* the number of variants, *h*^2^ the (SNP) heritability, ***γ*** the effect sizes on the allele scale, ***S*** the standard deviations of the genotypes, and ***β*** the effects of the scaled genotypes (Privé *et al*., 2020b). In LDpred2-auto, *p* and *h*^2^ are directly estimated within the Gibbs sampler, as opposed to testing several values of *p* and *h*^2^ from a grid of hyper-parameters (as in LDpred2-grid). This makes LDpred2-auto a method free of hyper-parameters which can therefore be applied directly without requiring a validation dataset to choose best-performing hyper-parameters (Privé *et al*., 2020b). *h*^2^ is estimated by ***β****^T^* ***Rβ***, where ***R*** is the correlation matrix between variants and ***β*** is a vector of causal effect sizes (after scaling) from one iteration of the Gibbs sampler. As for *p*, it is sampled from Beta(1 + *M_c_,* 1 + *M − M_c_*), where *M_c_* = *_j_* (*β_j_ /*= 0). Note a small change, we now sample *p* from Beta(1 + *M_c_/l*^-2^, 1 + (*M − M_c_*)*/l*^-2^), where *l*^-2^ is the average LD score, to add more variability in the sampling in order to account for a reduced effective number of (independent) variants.

Here we provide a new model and sampling scheme for LDpred2-auto that can be optionally used by setting parameter `use_MLE = TRUE` (otherwise it is run as described in the previous paragraph when using `use_MLE = FALSE`). In this new option, we extend LDpred2-auto with a third parameter *α* that controls the relationship between minor allele frequencies (or equivalently, standard deviations) of genotypes and expected effect sizes; the model becomes

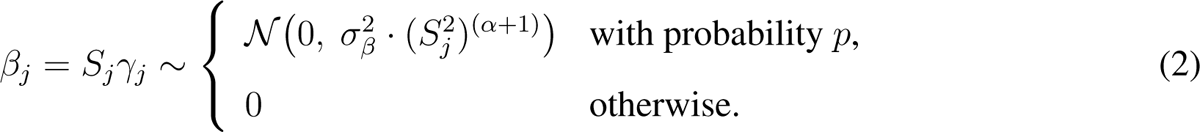

Therefore, it was earlier assumed that *α* = *−*1 and *σ*^2^ = *h*^2^/(*Mp*) in equation (1). This new model in equation (2) is similar to the model assumed by SBayesS, where *α* is denoted by *S* instead (Zeng *et al*., 2021). In SBayesS, *α* and *σ*^2^ are estimated by maximizing the likelihood of the normal distribution (over the causal variants from the Gibbs sampler). In the new LDpred2-auto, to add some sampling to these two parameters, we first sample causal variants with replacement before computing the maximum likelihood estimators. This maximum likelihood estimation (MLE) is implemented using R package roptim (Pan and Pan, 2022), and we bound the estimate of *α* to be between −1.5 and 0.5 (the default, but can be modified), and the estimate of *σ*^2^ to be between 0.5 and 2 times the one from the previous iteration of the Gibbs sampler. Note that we still estimate *h*^2^ = ***β****^T^* ***Rβ***, but that *h*^2^ is not used in the variance of sampled effect sizes anymore (Equation (2)). Note that this *h*^2^ estimation can be restricted to e.g. variants from a single LD block to get local heritability estimates. Finally, in both models and sampling schemes now implemented in LDpred2-auto, we now detect strong divergence when ***β****^T^* ***β*** *>* 2 **^^^***^T^* **^^^**, where ***β*** is the vector of scaled effect sizes from one iteration of the Gibbs sampler and ***β*^^^** is the marginal scaled effect sizes; corresponding chains are stopped and missing values are returned for effect sizes and estimates of missing iterations.

### Inference of predictive performance ***r*^2^**

To infer the out-of-sample predictive performance *r*^2^ (and its CI) of the resulting PGS from LDpred2-auto, we use the distribution of ***β*_1_***^T^* ***Rβ*_2_**, where ***β*_1_** and ***β*_2_** are two sampled vectors of causal effect sizes (after scaling) from two different chains of the Gibbs sampler. Intuitively, if prediction is perfect then ***β*_1_** = ***β*_2_** and *r*^2^ = *h*^2^; when power is very low, ***β*_1_** and ***β*_2_** are almost uncorrelated and *r*^2^ *≈* 0. Others have proposed to estimate *r*^2^ from a reference genotype set (Mak *et al*., 2017) or from an additional set of external GWAS summary statistics and LD reference (Pattee and Pan, 2020; Witteveen *et al*., 2022); here we only use the summary statistics that we input to LDpred2-auto. These previous works have shown that *r*^2^ can be approximated by (β-T β^)2/(β-T Rβ-) where ***β*^-^** and ***β*^^^** are respectively the predictive effects from the training set and the marginal effects from the test set (after scaling). Note that *E*[***β*_1_**] = ***β*^-^**, *E*[***β*** *^T^* ***Rβ***] = ***β*^-^***^T^****Rβ*^-^**, and, when the test sample has the same genetic ancestry as the GWAS used for training (to get the summary statistics), *E*[***Rβ*_2_**] *≈*. Therefore, we propose to use ***β*_1_***^T^* ***Rβ*_2_** as a sample of *r*^2^. This can also be computed for a specific chain by taking two samples ***β*_1_** and ***β*_2_** that are far enough on the same chain to remove the possible autocorrelation. This is what we use for SBayesS here, and also as an alternative means for post-filtering chains for prediction (cf. next Methods section).

In this paper, we check this new *r*^2^ approximation using extensive simulations (across many genetic architectures) and real data analyses (across 248 different phenotypes). These are compared to the partialr^2^ (on individual-level data from a separate test set). The partial correlation is computed with function `pcor` from R package bigstatsr, adjusting for the same eight covariates as in the GWAS, then squared (while keeping the sign). Corresponding 95% confidence intervals are estimated through bootstrapping individual indices.

### Post-filtering of chains in LDpred2-auto

Because a Gibbs sampler can be unstable, with so many variants and with possible mismatches between e.g. the GWAS summary statistics and the LD reference used, we have always recommended to run multiple chains in LDpred2-auto, and post-filter some of them as quality control (Privé *et al*., 2020b). We originally proposed to filter chains by keeping the ones providing PGS with the largest variances. Then we tested an almost equivalent and simpler alternative (Privé *et al*., 2022a), which is to keep only chains that provide top imputed marginal scaled effect sizes *β*^̌^ = *Rβ*^-^, where *R* is the correlation matrix and *β*^-^ are the PGS scaled effect sizes. This is the default post-filtering of LDpred2-auto chains that we use here, for both prediction and inference.

Moreover, here we test two other filtering criteria in the first simulations based on continuous outcomes (and call this “LDpred2_newfilter”). First, for prediction, we test filtering on the average of ***β*_1_***^T^* ***Rβ*_2_** within each chain, which is an estimate of *r*^2^ (cf. previous Methods section). Second, for inference, we filter on some convergence criterion. The split-Rhat statistic is a popular metric to test for good mixing and convergence of Markov chains (Vehtari *et al*., 2021). However, we have found this statistic to perform badly when e.g. one parameter gets stuck and is constant; in this case, the chain does not mix well, but a perfect Rhat of 1.0 is obtained. Instead, we have found that a two-sample Cramer-von Mises statistic (Anderson, 1962), by similarly using both parts of the chain after burn-in, is often highly correlated with the split-Rhat statistic while not suffering from the previous issue. We therefore chose to use this statistic, and average the three statistics computed for *h*^2^, *p*, and *α* for each chain. We use a threshold of 4 above which we filter out the chain, because we have found that a value of 4 for this statistic often corresponds to a value close to 1.05 for the split-Rhat.

## Acknowledgements

Authors thank members of the NCRR/QGG StatGen group and Marc-André Legault for helpful discussions, as well as reviewers for their useful comments. Authors also thank GenomeDK and Aarhus University for providing computational resources and support that contributed to these research results. This research has been conducted using the UK Biobank Resource under Application Number 58024; authors thank all the UK Biobank participants for contributing to such useful data for Research.

## Funding

F.P., C.A. and B.J.V. are supported by the Danish National Research Foundation (Niels Bohr Professorship to Prof. John McGrath), the Lundbeck Foundation Initiative for Integrative Psychiatric Research, iPSYCH (R102-A9118, R155-2014-1724, R248-2017-2003), and a Lundbeck Foundation Fellowship (R335-2019-2339 to B.J.V.).

## Declaration of Interests

B.J.V. is on Allelica’s international advisory board. The other authors have no competing interests to declare.

## Code and data availability

The UK Biobank data is available through a procedure described at https://www.ukbiobank.ac.uk/using-the-resource/. Descriptions of UK Biobank phenotypes used here are available at https://github.com/privefl/paper-infer/blob/main/phenotype-description.tsv. LD matrices for HapMap3+ variants computed from the N.W. European UKBB data used in this paper are available at https://doi.org/10.6084/m9.figshare.21305061. All code used for this paper is available at https://github.com/privefl/paper-infer/tree/master/code. We have extensively used R packages bigstatsr and bigsnpr (Privé *et al*., 2018) for analyzing large genetic data, packages from the future framework (Bengtsson, 2021) for easy scheduling and parallelization of analyses on the HPC cluster, and packages from the tidyverse suite (Wickham *et al*., 2019) for shaping and visualizing results. The latest version of R package bigsnpr can be installed from GitHub, and a recent enough version can be installed from CRAN.

## Notes

### Summary of Updates

Added more analyses + clarified a few points.

