## Supplementary Materials for "Inferring disease architecture and predictive ability with LDpred2-auto"

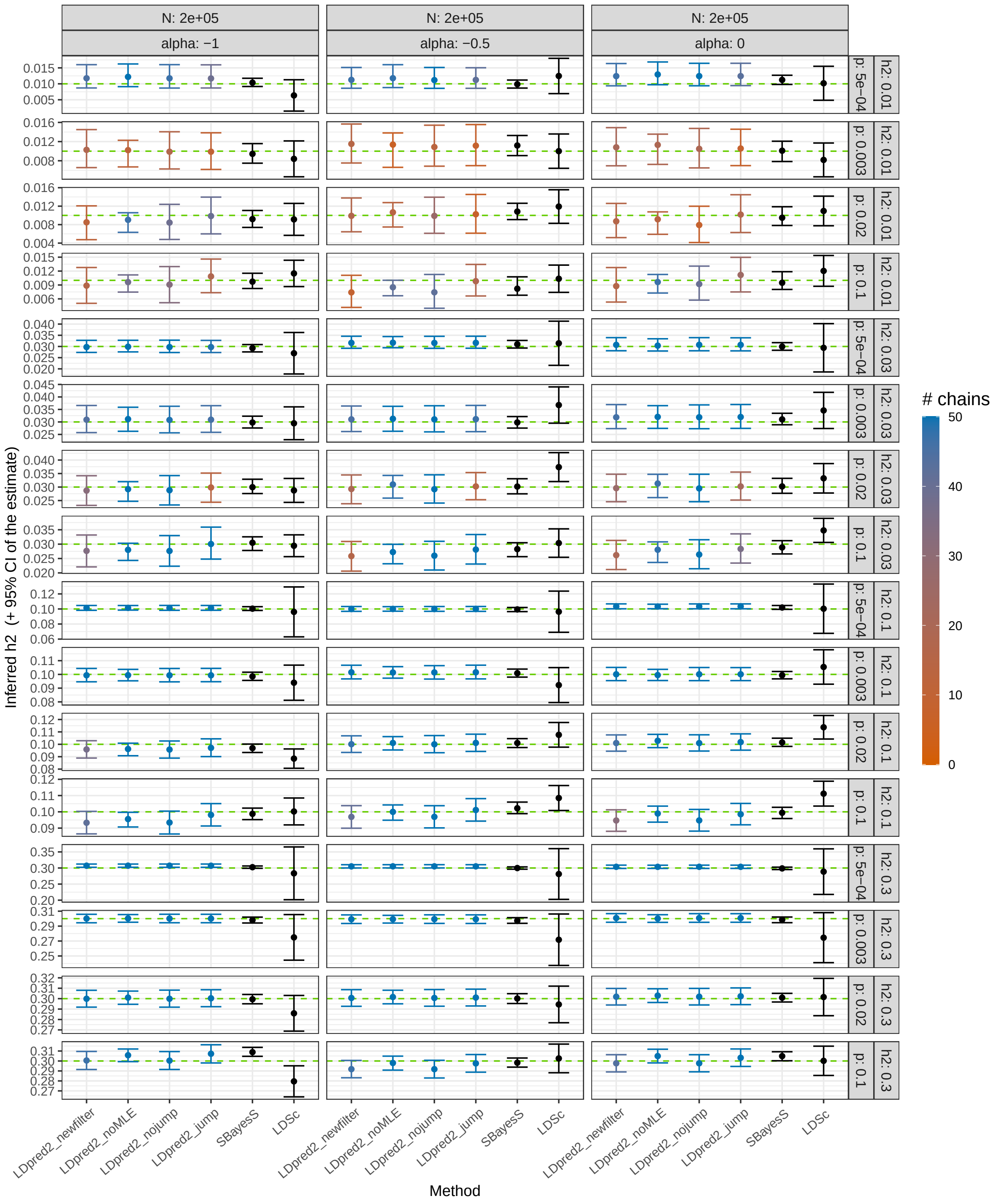


Figure S1: Inferred SNP heritability $h^2$ in simulations with continuous outcomes and N=200K. Horizontal dashed lines represent the true simulated values. For LDpred2-auto, suffix "nojump"/"jump" refers to using `allow_jump_sign = FALSE/TRUE` (and `use_MLE = TRUE`), and "noMLE" refers to using `use_MLE = FALSE` (and `allow_jump_sign = FALSE`), and "newfilter" is similar to "nojump" but uses a different post-filtering of chains (Methods). Note that the recommended option is to use `allow_jump_sign = FALSE`[4]. The 95% confidence intervals for the LDpred2-auto and SBayesS estimates are obtained from the 2.5% and 97.5% quantiles of all the $h^2$ estimates from the iterations (after burn-in) of the chains kept (note that only one chain is used and kept in SBayesS). The 95% confidence interval for the LD Score (LDSc) regression estimate is obtained as ±1.96 of its standard error. Colors for LDpred2-auto models represent the number of chains kept (out of 50).

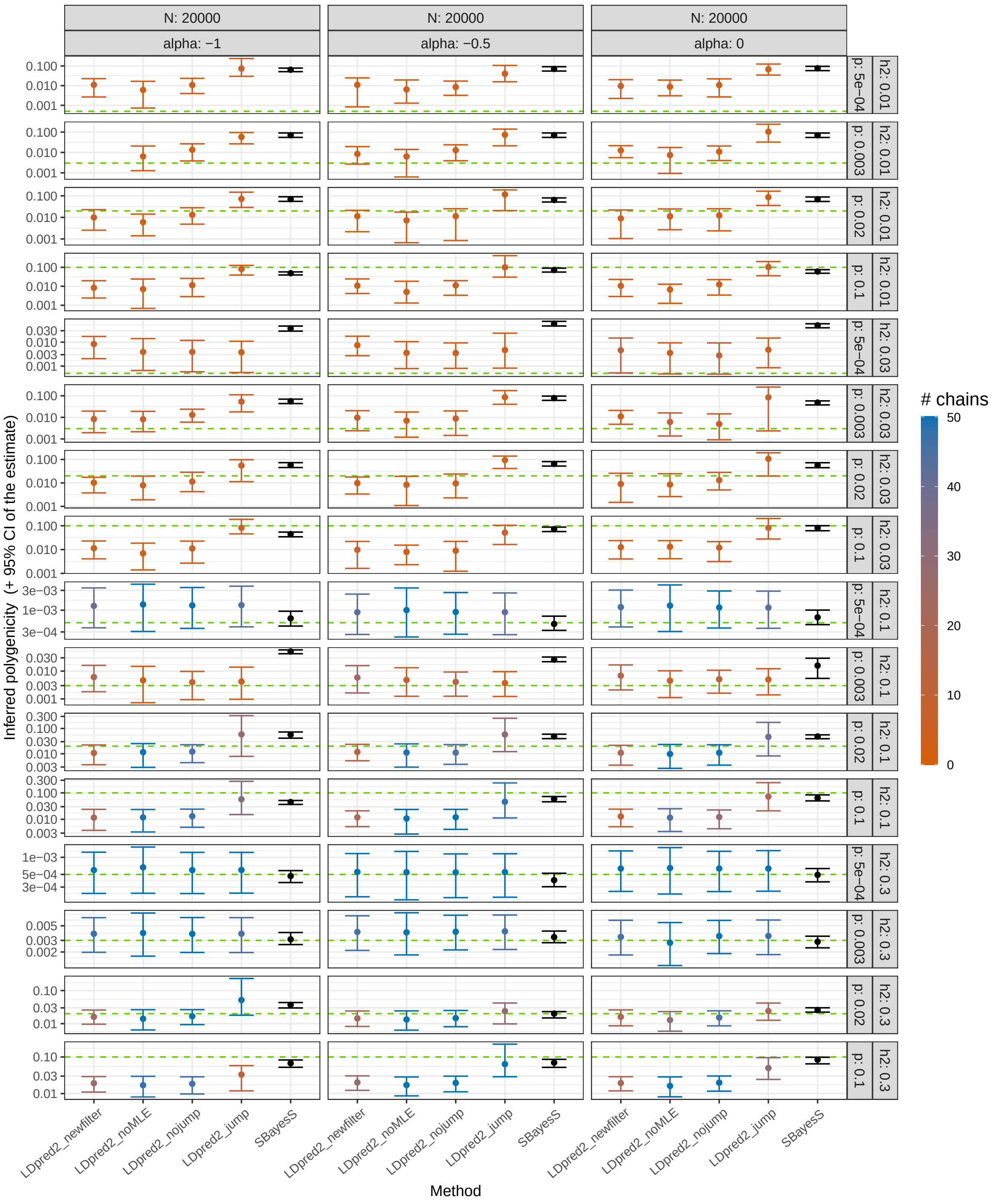


Figure S2: Inferred polygenicity $p$ in simulations with continuous outcomes and N=20K. Horizontal dashed lines represent the true simulated values. For LDpred2-auto, suffix "nojump"/"jump" refers to using `allow_jump_sign = FALSE/TRUE` (and `use_MLE = TRUE`), and "noMLE" refers to using `use_MLE = FALSE` (and `allow_jump_sign = FALSE`), and "newfilter" is similar to "nojump" but uses a different post-filtering of chains (Methods). Note that the recommended option is to use `allow_jump_sign = FALSE`[4]. The 95% confidence intervals for the LDpred2-auto and SBayesS estimates are obtained from the 2.5% and 97.5% quantiles of all the $p$ estimates from the iterations (after burn-in) of the chains kept (note that only one chain is used and kept in SBayesS). Colors for LDpred2-auto models represent the number of chains kept (out of 50).

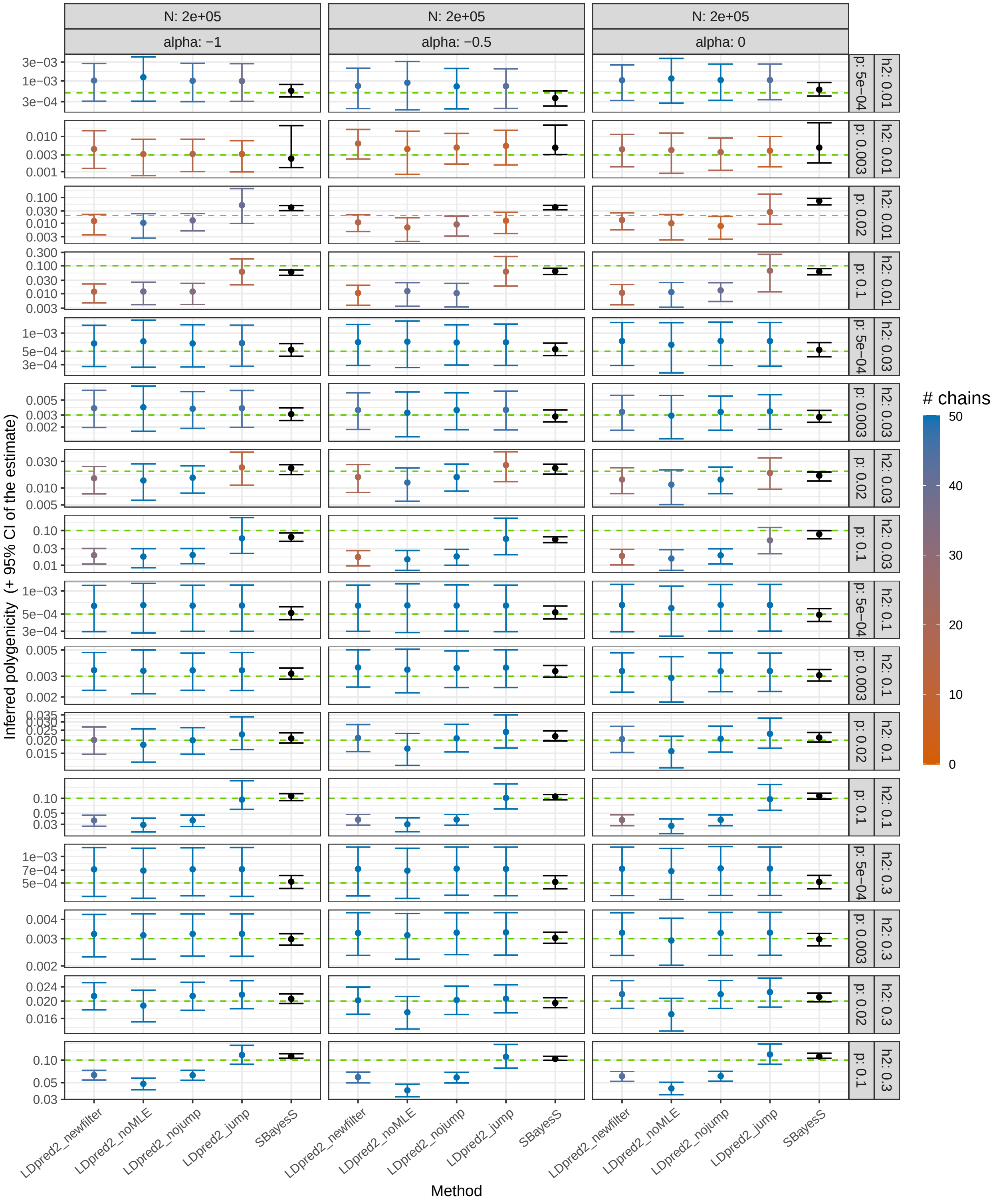


Figure S3: Inferred polygenicity $p$ in simulations with continuous outcomes and N=200K. Horizontal dashed lines represent the true simulated values. For LDpred2-auto, suffix "nojump"/"jump" refers to using `allow_jump_sign = FALSE/TRUE` (and `use_MLE = TRUE`), and "noMLE" refers to using `use_MLE = FALSE` (and `allow_jump_sign = FALSE`), and "newfilter" is similar to "nojump" but uses a different post-filtering of chains (Methods). Note that the recommended option is to use `allow_jump_sign = FALSE`[4]. The 95% confidence intervals for the LDpred2-auto and SBayesS estimates are obtained from the 2.5% and 97.5% quantiles of all the $p$ estimates from the iterations (after burn-in) of the chains kept (note that only one chain is used and kept in SBayesS). Colors for LDpred2-auto models represent the number of chains kept (out of 50).

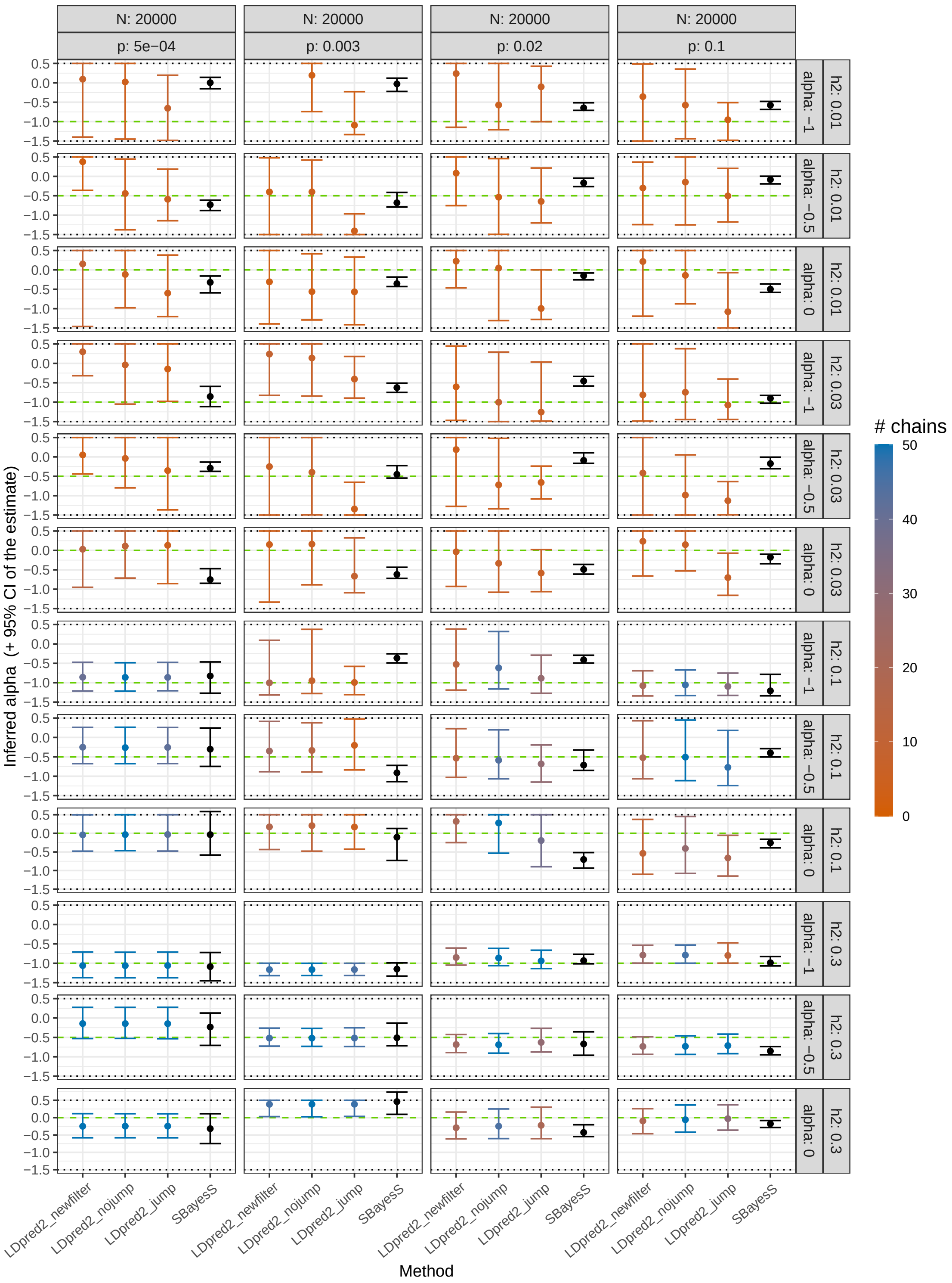


Figure S4: Inferred $\alpha$ in simulations with continuous outcomes and N=20K. Horizontal dashed lines represent the true simulated values. Horizontal dotted lines represent boundaries imposed on the LDpred2-auto estimates. For LDpred2-auto, suffix "nojump"/"jump" refers to using `allow_jump_sign = FALSE/TRUE` (and `use_MLE = TRUE`), and "newfilter" is similar to "nojump" but uses a different post-filtering of chains (Methods). Note that "LDpred2_noMLE" (`use_MLE = FALSE`) does not infer $\alpha$. Note that the recommended option is to use `allow_jump_sign = FALSE`[4]. The 95% confidence intervals for the LDpred2-auto and SBayesS estimates are obtained from the 2.5% and 97.5% quantiles of all the $\alpha$ estimates from the iterations (after burn-in) of the chains kept (note that only one chain is used and kept in SBayesS). Colors for LDpred2-auto models represent the number of chains kept (out of 50).

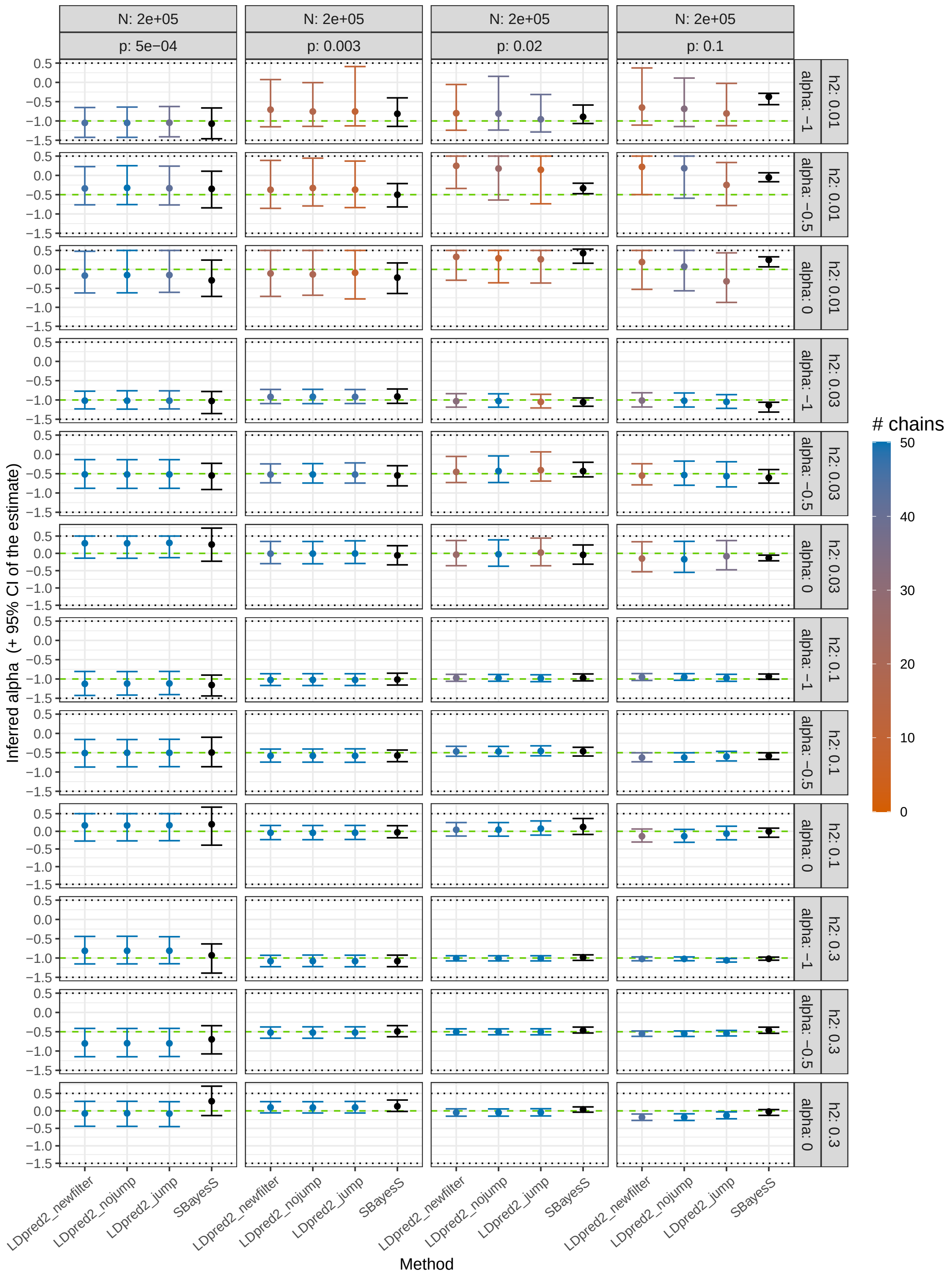


Figure S5: Inferred $\alpha$ in simulations with continuous outcomes and N=200K. Horizontal dashed lines represent the true simulated values. Horizontal dotted lines represent boundaries imposed on the LDpred2-auto estimates. For LDpred2-auto, suffix "nojump"/"jump" refers to using `allow_jump_sign = FALSE/TRUE` (and `use_MLE = TRUE`), and "newfilter" is similar to "nojump" but uses a different post-filtering of chains (Methods). Note that "LDpred2_noMLE" (`use_MLE = FALSE`) does not infer $\alpha$. Note that the recommended option is to use `allow_jump_sign = FALSE`[4]. The 95% confidence intervals for the LDpred2-auto and SBayesS estimates are obtained from the 2.5% and 97.5% quantiles of all the $\alpha$ estimates from the iterations (after burn-in) of the chains kept (note that only one chain is used and kept in SBayesS). Colors for LDpred2-auto models represent the number of chains kept (out of 50).

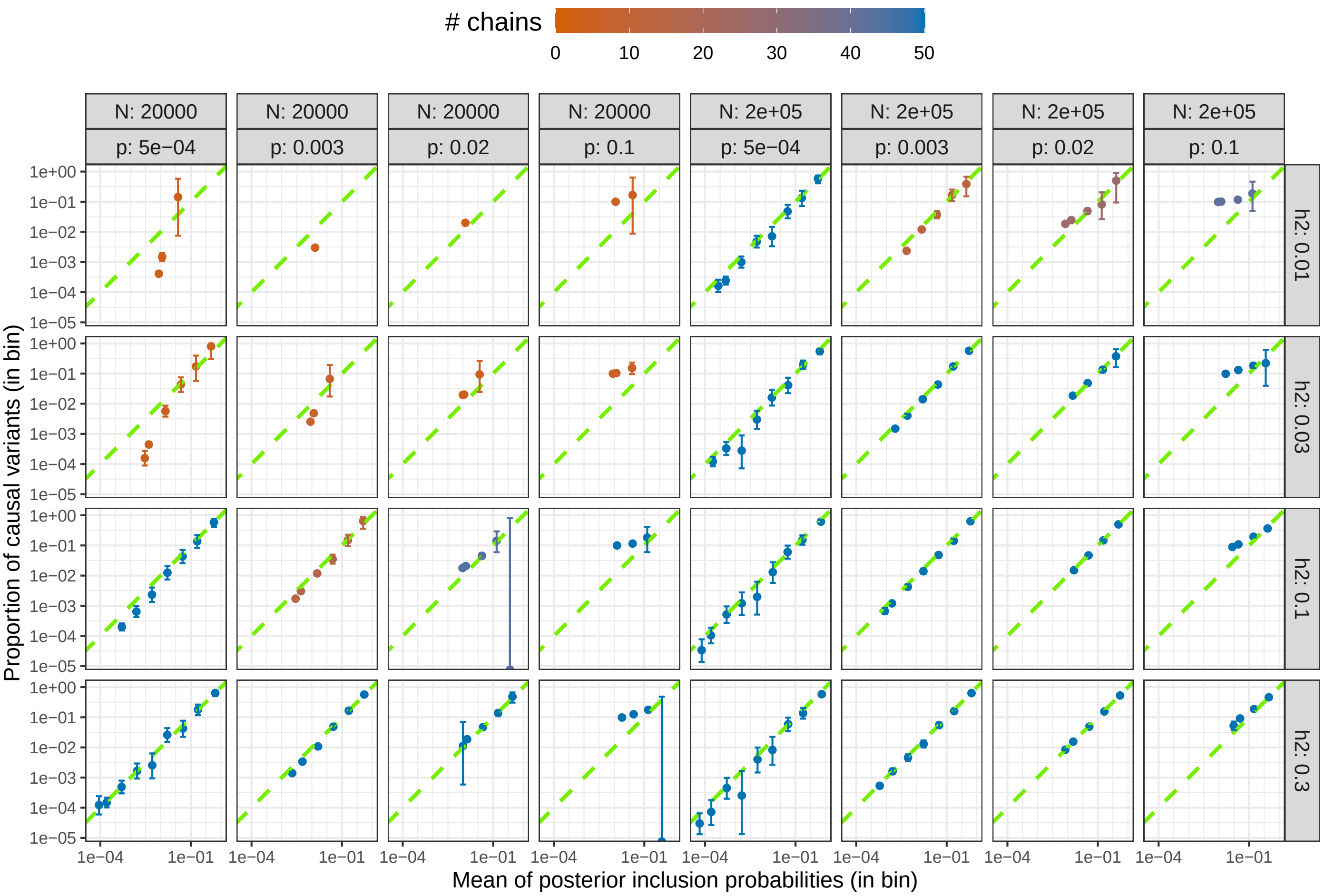


Figure S6: LDpred2-auto calibration of per-variant posterior probabilities of being causal (also known as posterior inclusion probabilities) in simulations with continuous outcomes. These probabilities are binned on a log-scale, and mean in each bin is compared to the proportion of simulated causal variants in the bin (with 95% CI). Green dashed lines represent the 1:1 line. Only results for $\alpha = -0.5$ and `allow_jump_sign = FALSE` are represented. Colors represent the number of chains kept (out of 50).

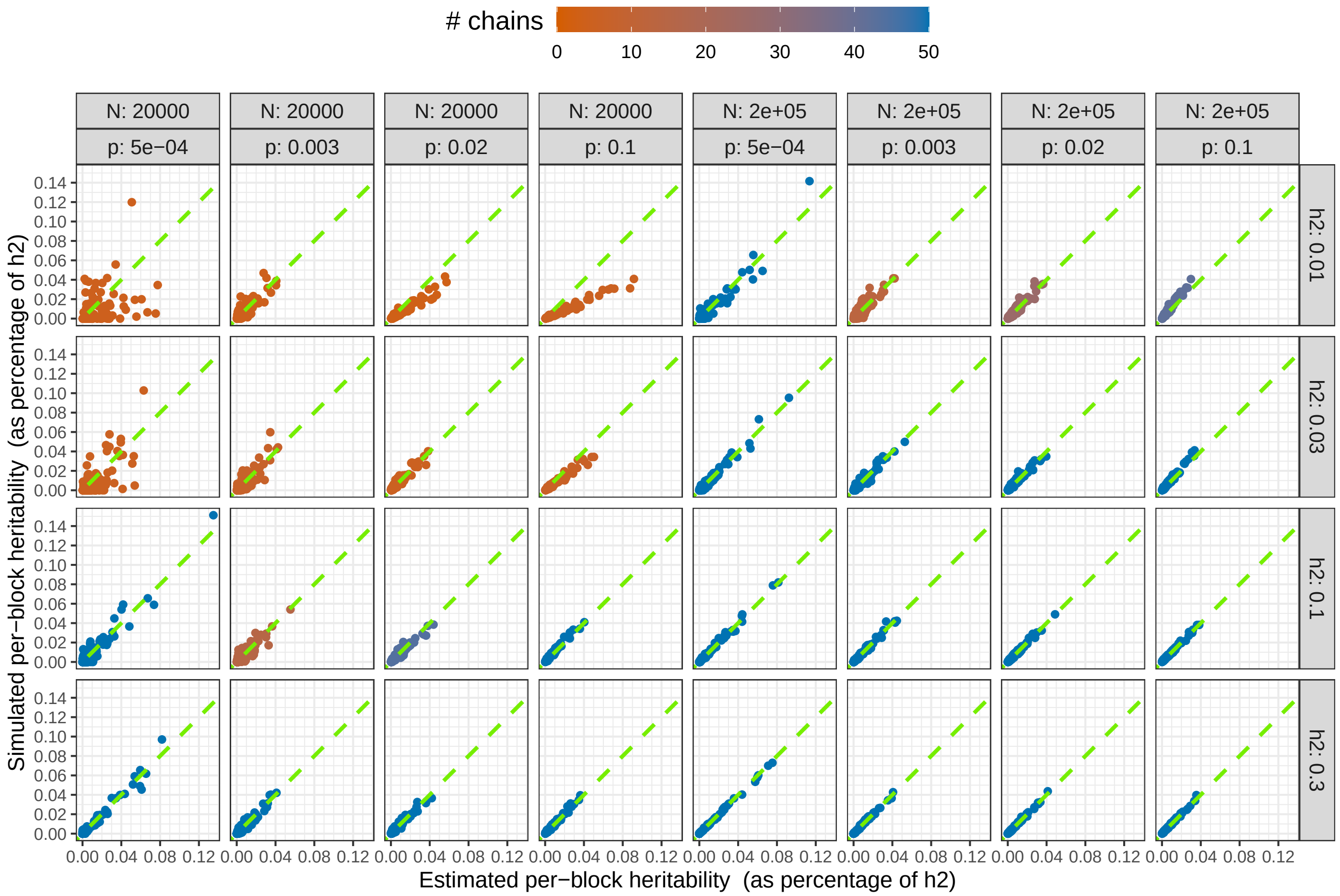


Figure S7: LDpred2-auto calibration of per-block heritability estimates in simulations with continuous outcomes. Local $h^2$ estimate for each block is compared to the true simulated local heritability in the block. Green dashed lines represent the 1:1 line. Only results for $\alpha = -0.5$ and `allow_jump_sign = FALSE` are represented. Colors represent the number of chains kept (out of 50).

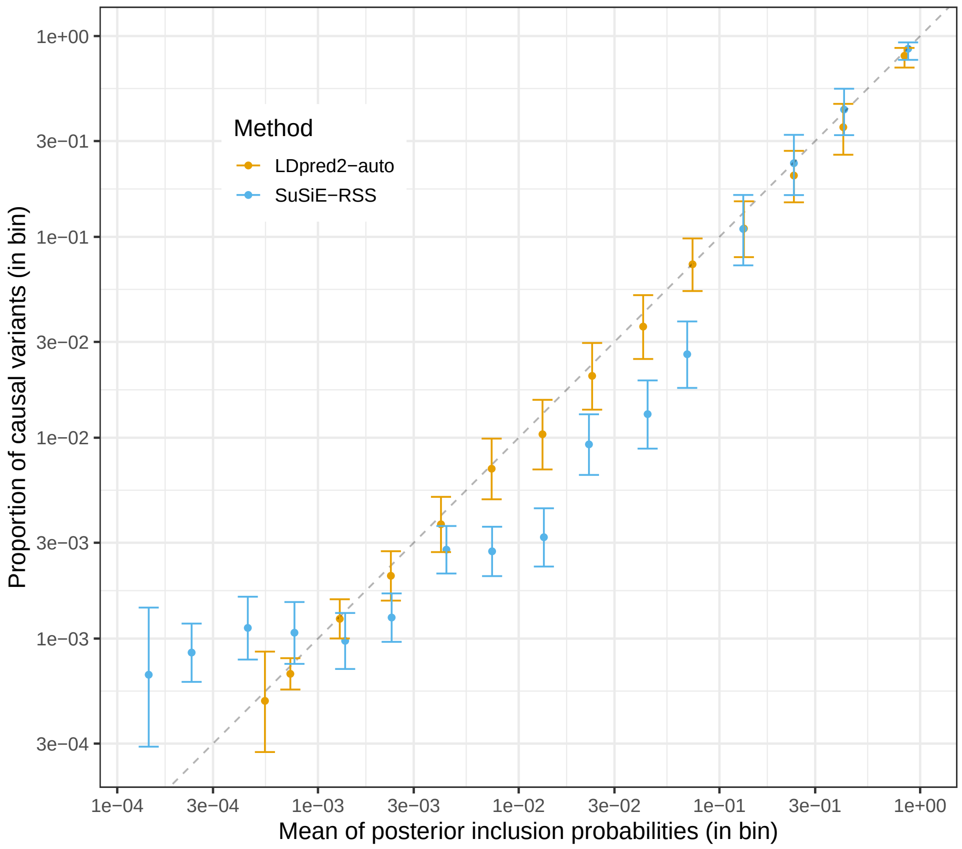


(a) With $p = 0.002$.

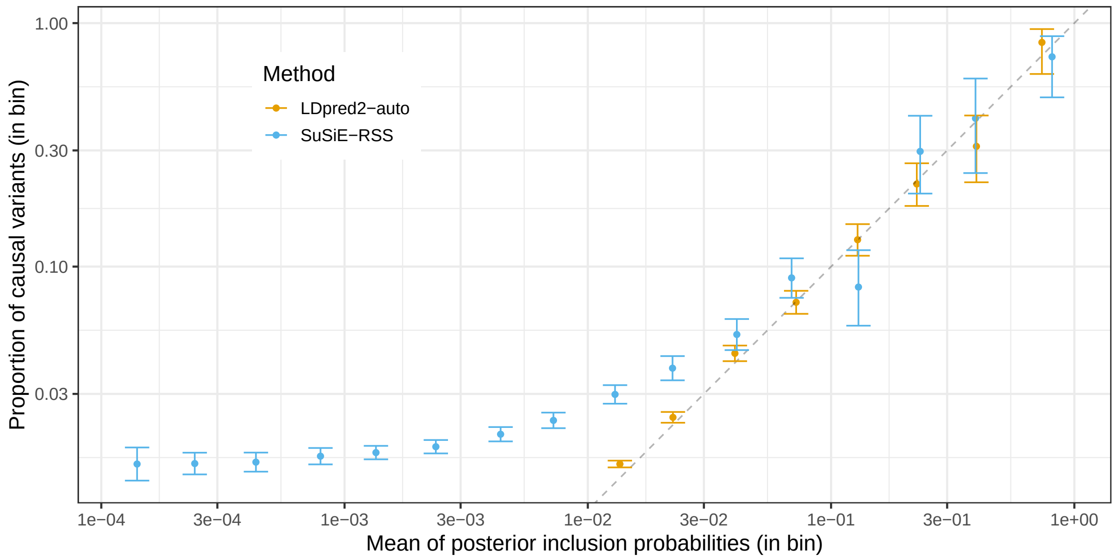


(b) With $p = 0.02$.

Figure S8: Calibration of per-variant posterior probabilities of being causal in two simulations with continuous outcomes, assuming $h^2 = 0.1$, $\alpha = -0.5$, and $N = 10^5$.

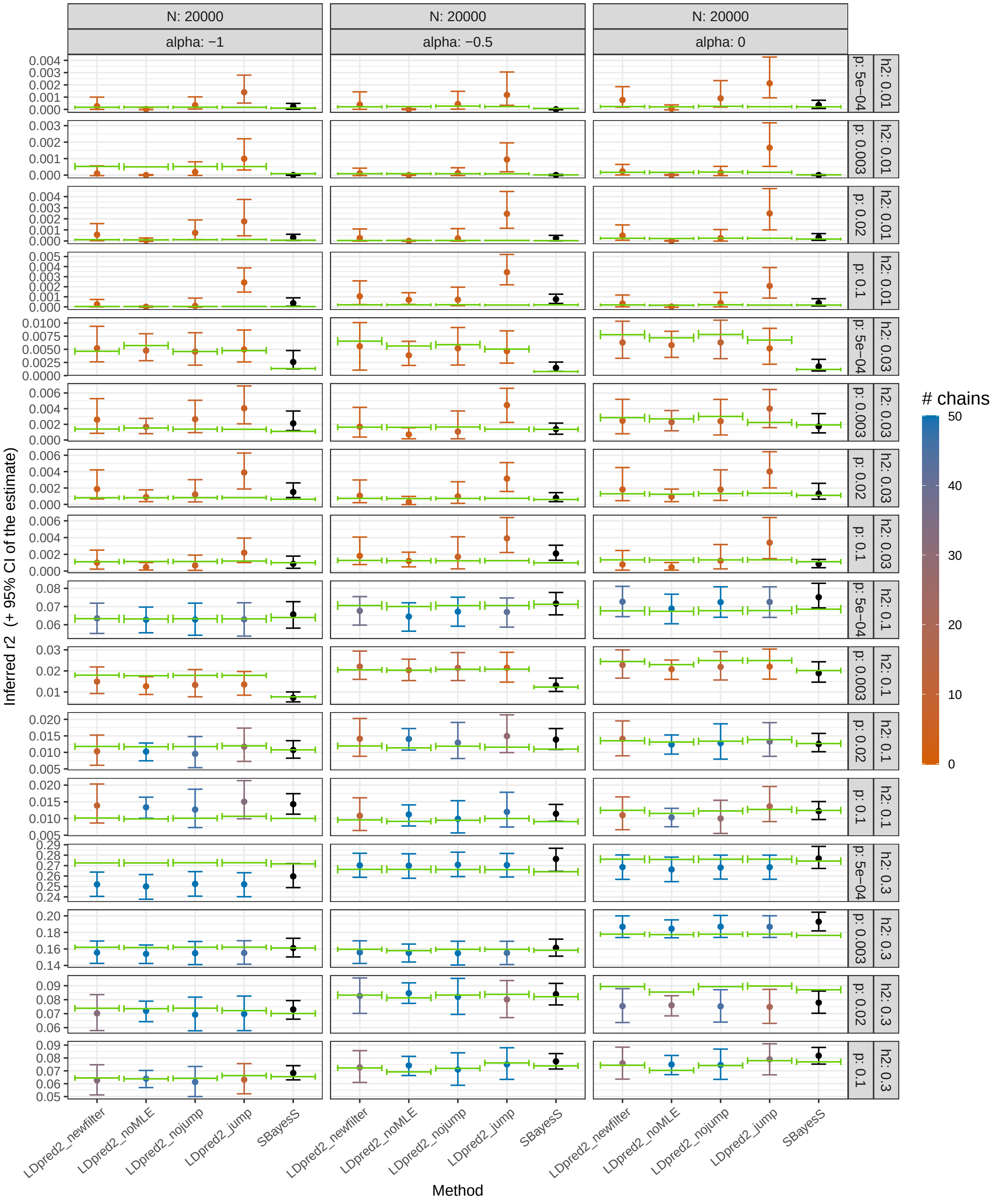


Figure S9: Inferred predictive performance $r^2$ in simulations with continuous outcomes and N=20K. Green segments represent $r^2$ in the test set. For LDpred2-auto, suffix "nojump"/"jump" refers to using `allow_jump_sign = FALSE/TRUE` (and `use_MLE = TRUE`), and "noMLE" refers to using `use_MLE = FALSE` (and `allow_jump_sign = FALSE`), and "newfilter" is similar to "nojump" but uses a different post-filtering of chains (Methods). Note that the recommended option is to use `allow_jump_sign = FALSE`[4]. Colors for LDpred2-auto models represent the number of chains kept (out of 50).

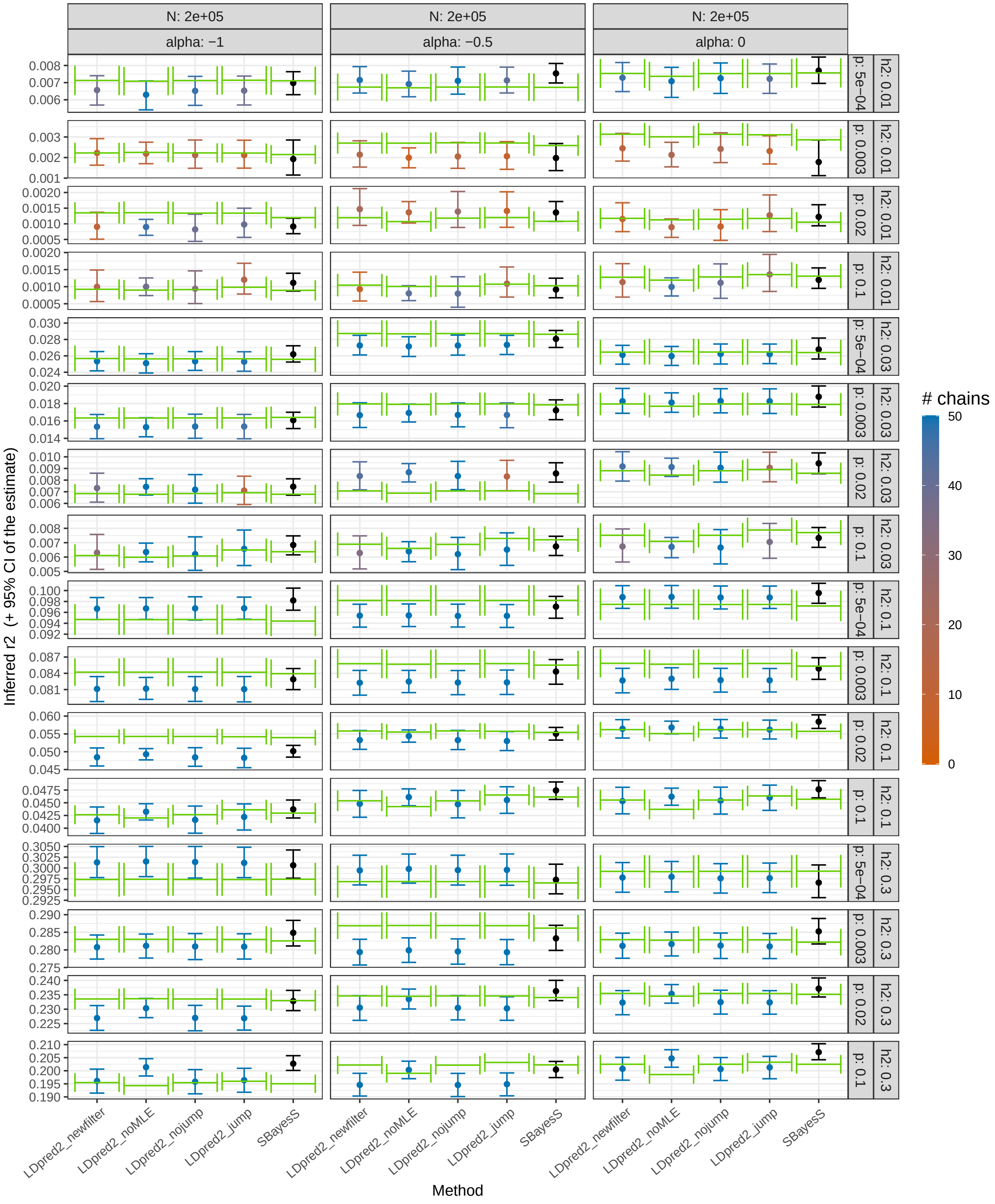


Figure S10: Inferred predictive performance $r^2$ in simulations with continuous outcomes and N=200K. Green segments represent $r^2$ in the test set. For LDpred2-auto, suffix "nojump"/"jump" refers to using `allow_jump_sign = FALSE/TRUE` (and `use_MLE = TRUE`), and "noMLE" refers to using `use_MLE = FALSE` (and `allow_jump_sign = FALSE`), and "newfilter" is similar to "nojump" but uses a different post-filtering of chains (Methods). Note that the recommended option is to use `allow_jump_sign = FALSE`[4]. Colors for LDpred2-auto models represent the number of chains kept (out of 50).

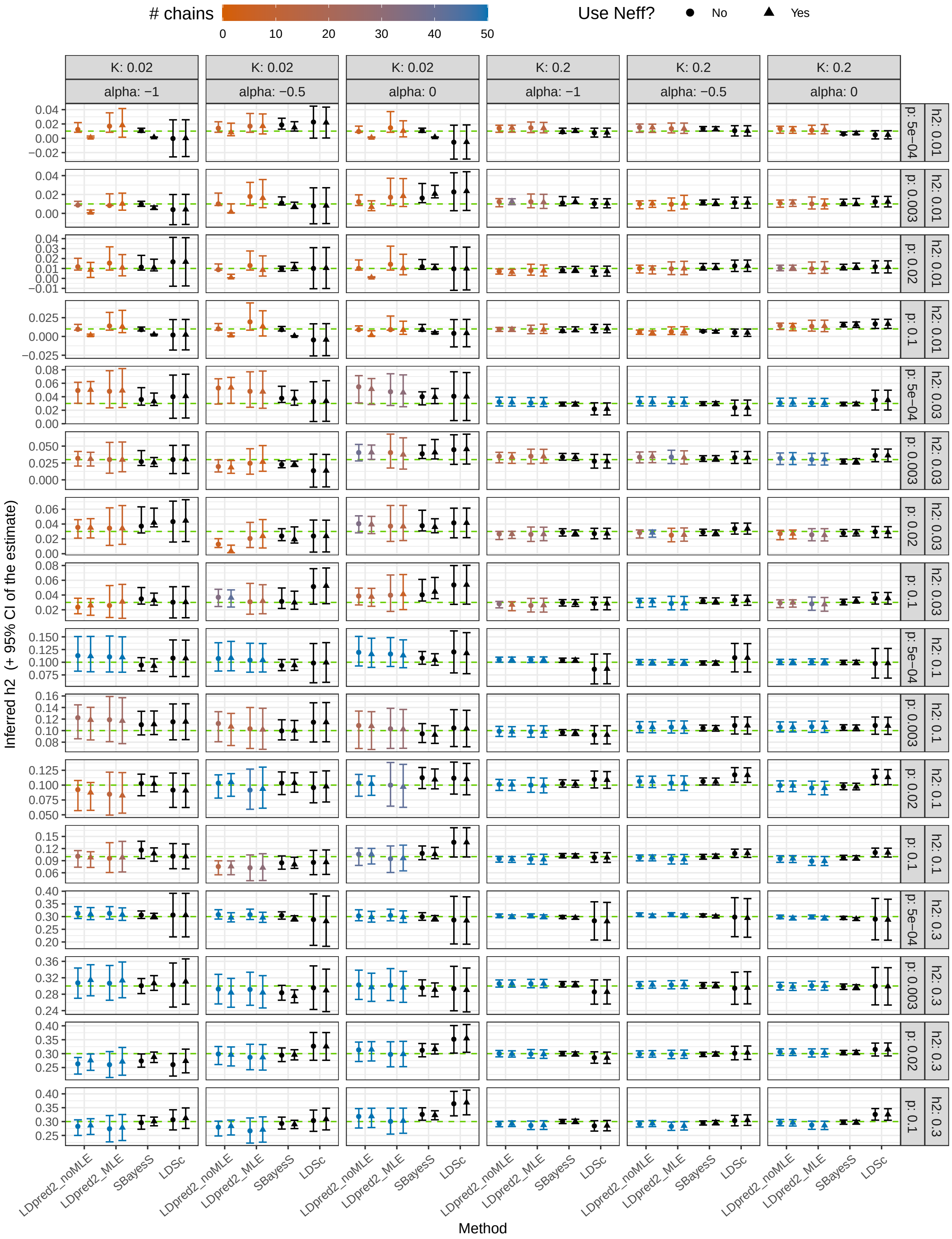


Figure S11: Inferred SNP heritability $h^2$ in simulations with binary outcomes. Horizontal dashed lines represent the true simulated values. The 95% confidence interval for the LDpred2-auto estimate is obtained from the 2.5% and 97.5% quantiles of all the $h^2$ estimates from the iterations (after burn-in) of the chains kept. The 95% confidence interval for the LD Score regression estimate is obtained from $\pm 1.96$ times its standard error. Colors for LDpred2-auto models represent the number of chains kept (out of 50). Option `Neff` controls whether a logistic regression is used for the GWAS with the effective sample in LDpred2, or a linear regression and then the total sample size. All $h^2$ estimates are transformed to the liability scale with K_pop=K and K_GWAS either K (the simulated prevalence) or 0.5 (when using Neff) using function `coef_to_liab` from R package bigsnpr[2,3].

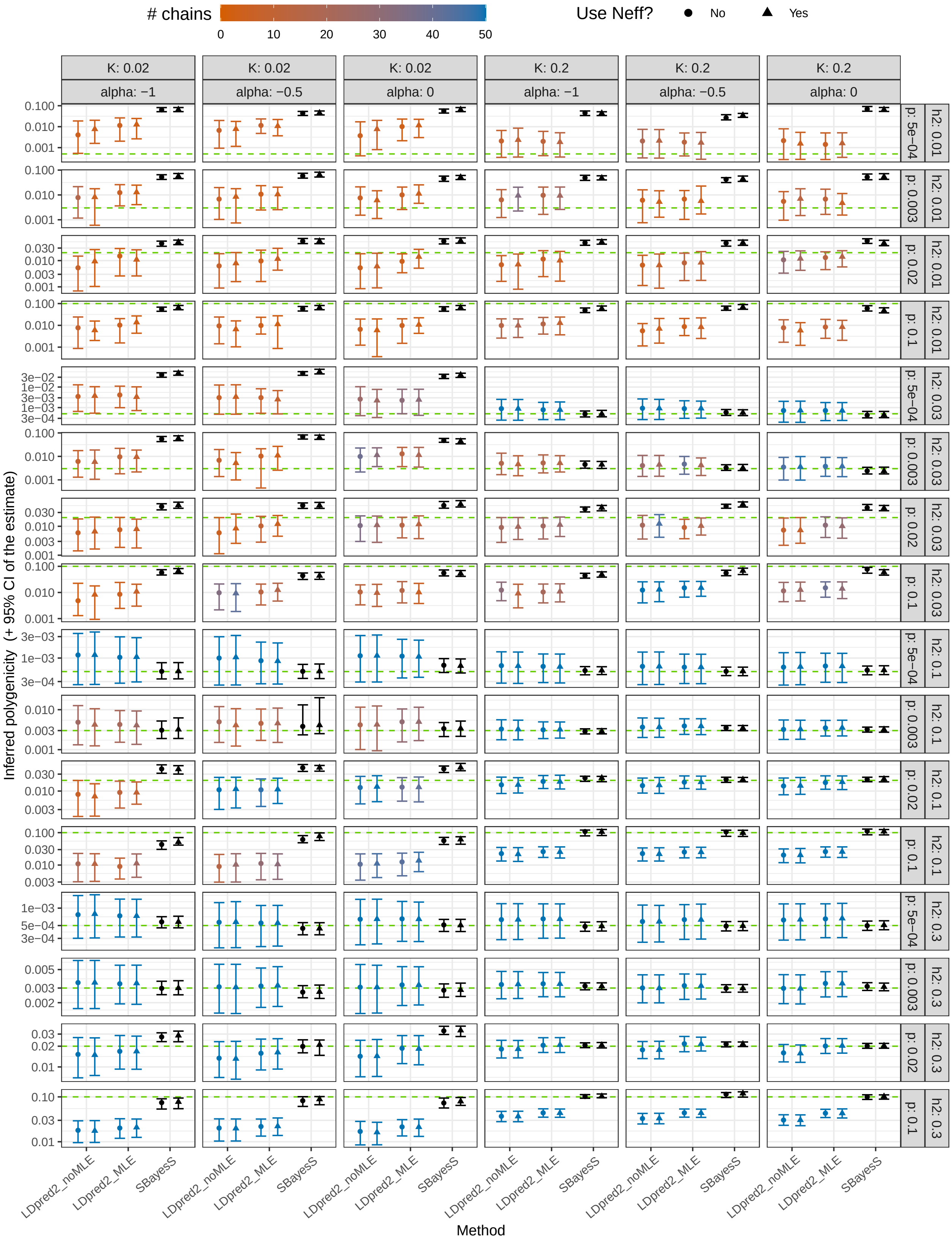


Figure S12: Inferred polygenicity $p$ in simulations with binary outcomes. Horizontal dashed lines represent the true simulated values. Colors for LDpred2-auto models represent the number of chains kept (out of 50). Option `Neff` controls whether a logistic regression is used for the GWAS with the effective sample in LDpred2, or a linear regression and then the total sample size.

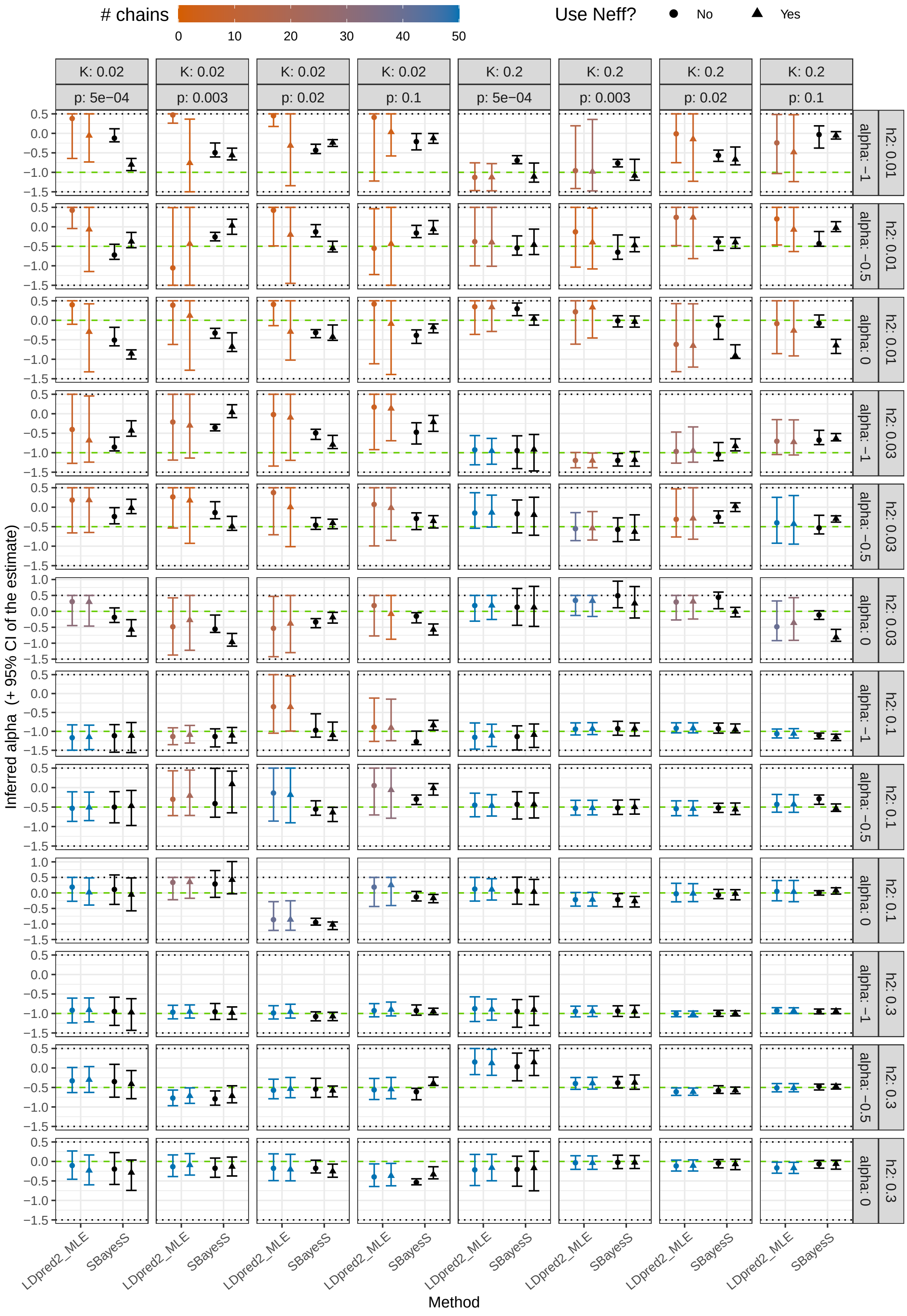


Figure S13: Inferred $\alpha$ in simulations with binary outcomes. Horizontal dashed lines represent the true simulated values. Horizontal dotted lines represent boundaries imposed on the estimates. Colors for LDpred2-auto models represent the number of chains kept (out of 50). Option `Neff` controls whether a logistic regression is used for the GWAS with the effective sample in LDpred2, or a linear regression and then the total sample size.

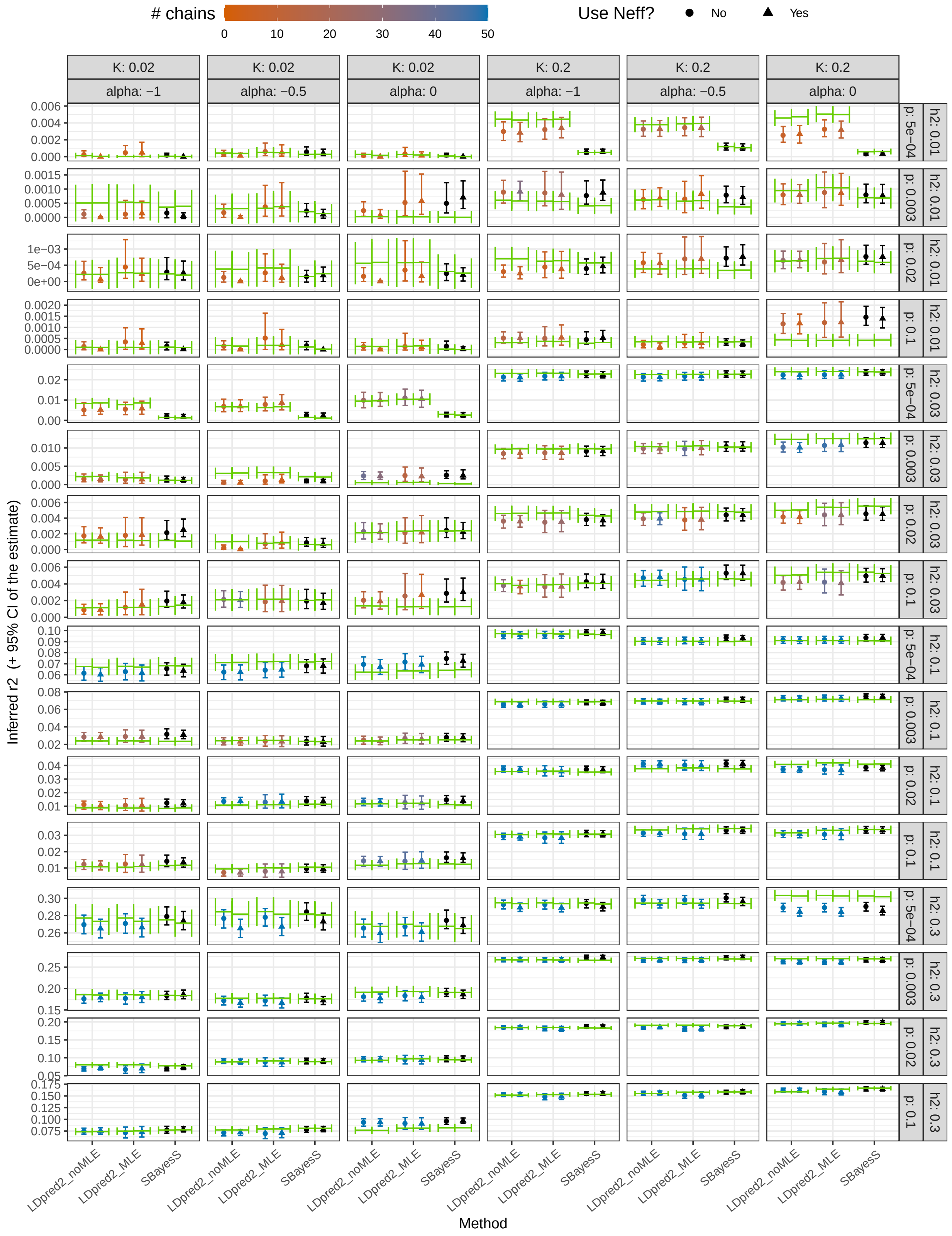

Figure S14: Inferred predictive performance $r^2$ in simulations with binary outcomes. Green segments represent $r^2$ in the test set. Colors for LDpred2-auto models represent the number of chains kept (out of 50). Option `Neff` controls whether a logistic regression is used for the GWAS with the effective sample in LDpred2, or a linear regression and then the total sample size. All $r^2$ estimates are transformed to the liability scale with K_pop=K and K_GWAS either K (the simulated prevalence) or 0.5 (when using Neff) using function `coef_to_liab` from R package bigsnpr[2,3]; the ones from the test set are transformed with K and K.

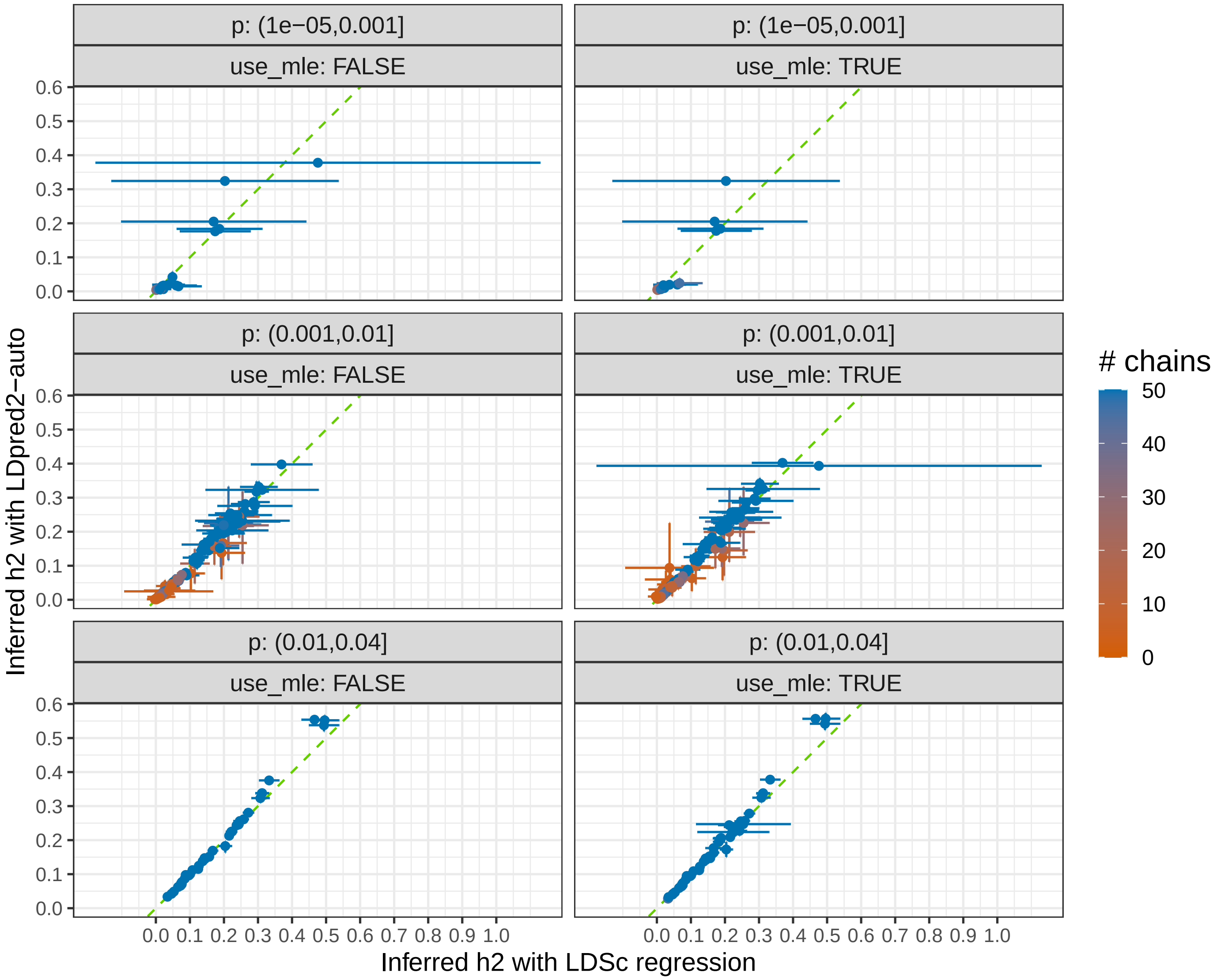


Figure S15: SNP heritability estimates from either LDpred2-auto or LD Score regression for all 248 phenotypes defined from the UK Biobank. These are stratified by the polygenicity estimated from LDpred2-auto. Green dashed lines represent the 1:1 line. The 95% confidence interval for the LDpred2-auto estimate is obtained from the 2.5% and 97.5% quantiles of all the $h^2$ estimates from the iterations (after burn-in) of the chains kept. The 95% confidence interval for the LD Score regression estimate is obtained from $\pm$1.96 times its standard error. “use_mle: TRUE” corresponds to using the new 3-parameter model and sampling scheme (Methods). Colors represent the number of chains kept for LDpred2-auto (out of 50).

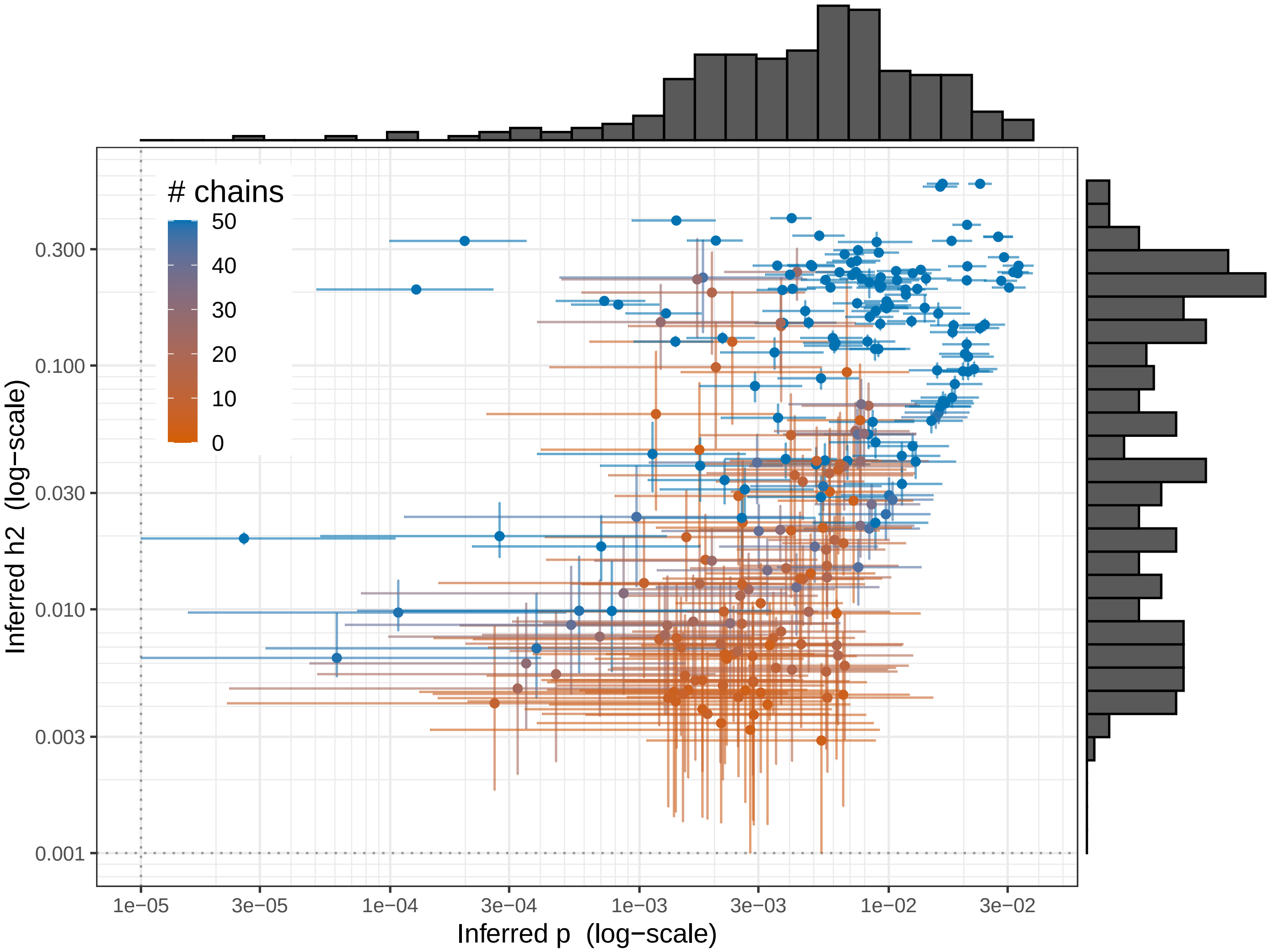


Figure S16: Estimates from LDpred2-auto for either the SNP heritability $h^2$ or the polygenicity $p$ for all 248 phenotypes defined from the UK Biobank. Estimates of $h^2$ are constrained to be at least 0.001, and at least $10^{-5}$ for $p$. The 95% confidence interval for the LDpred2-auto estimate is obtained from the 2.5% and 97.5% quantiles of all the estimates from the iterations (after burn-in) of the chains kept. Colors represent the number of chains kept for LDpred2-auto (out of 50).

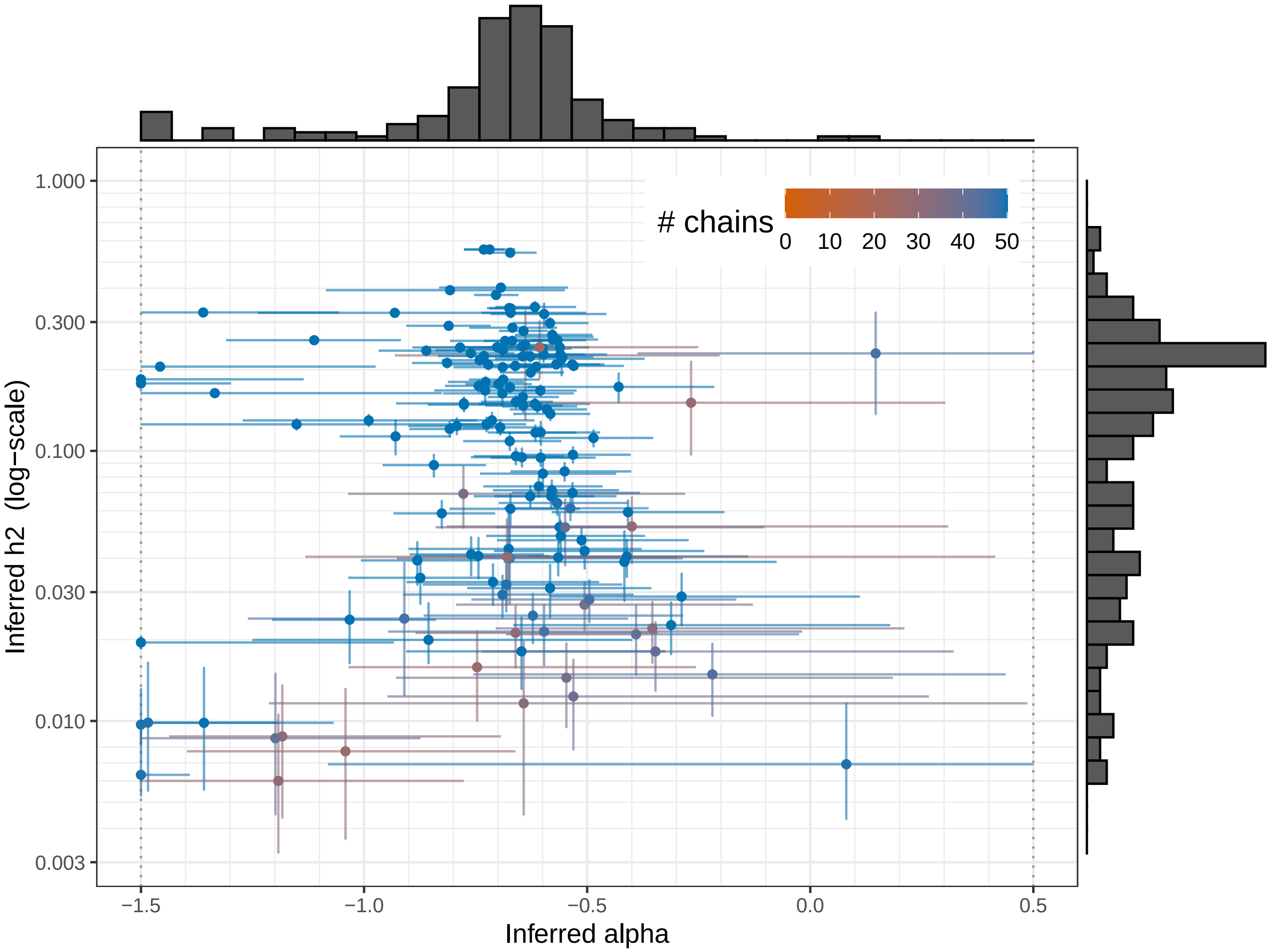


Figure S17: Estimates from LDpred2-auto for either the SNP heritability $h^2$ or $\alpha$ for all 248 phenotypes defined from the UK Biobank. The 95% confidence interval for the LDpred2-auto estimate is obtained from the 2.5% and 97.5% quantiles of all the estimates from the iterations (after burn-in) of the chains kept. Colors represent the number of chains kept for LDpred2-auto (out of 50). We only show phenotypes for which there are more than 25 chains kept, because simulations have shown that $\alpha$ estimates are unreliable when a small number of chains is kept (Figure S4).

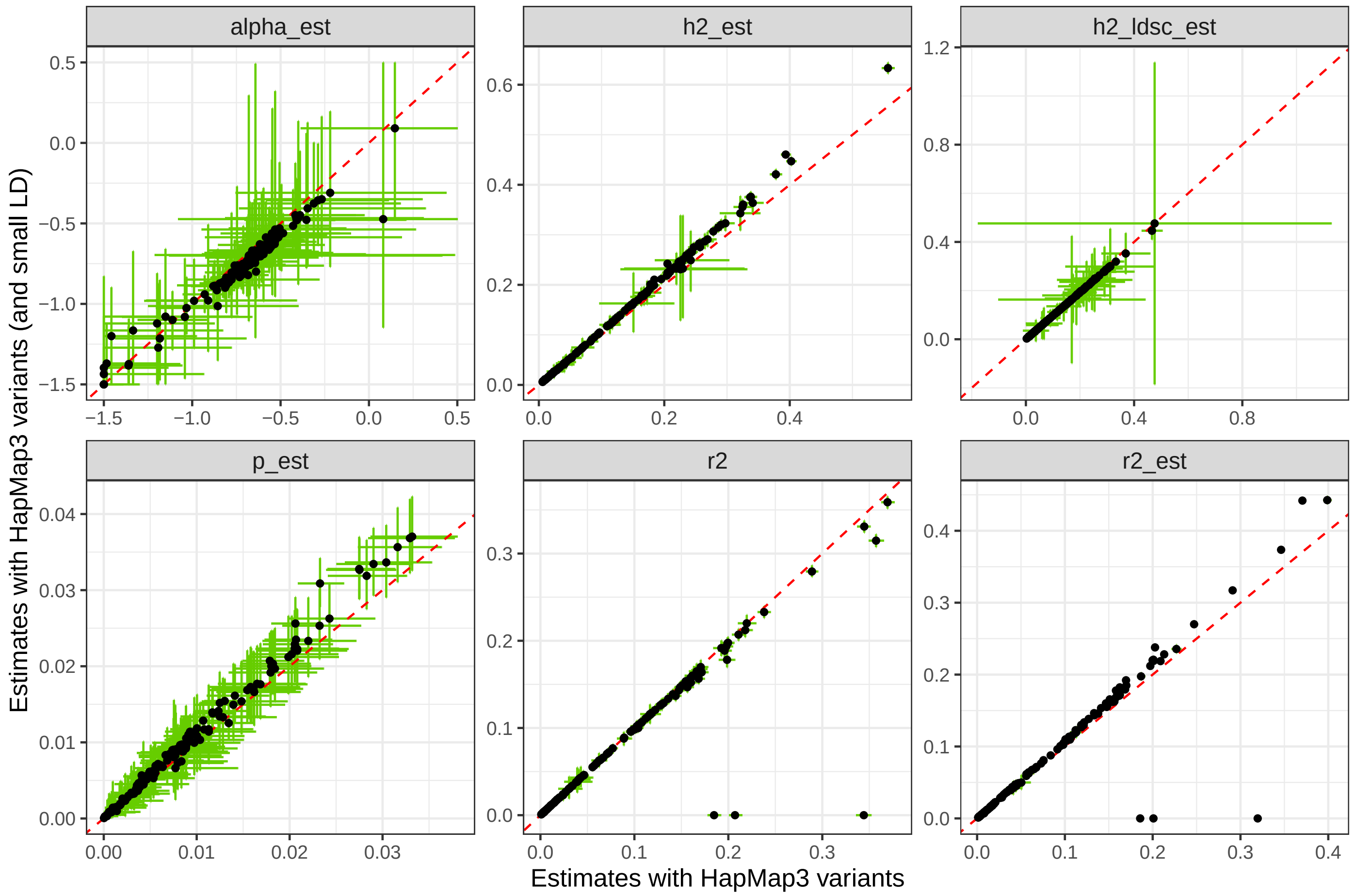


Figure S18: LDpred2-auto estimates for UKBB phenotypes with either a small or a large LD reference. Only 154 phenotypes with more than 25 chains kept when using the large LD reference are represented here. Red dashed lines represent the 1:1 line. The 95% confidence interval for the LDpred2-auto estimate (in green) is obtained from the 2.5% and 97.5% quantiles of all the estimates from the iterations (after burn-in) of the chains kept. The 95% confidence interval for $r^2$ in the test set is obtained from bootstrap.

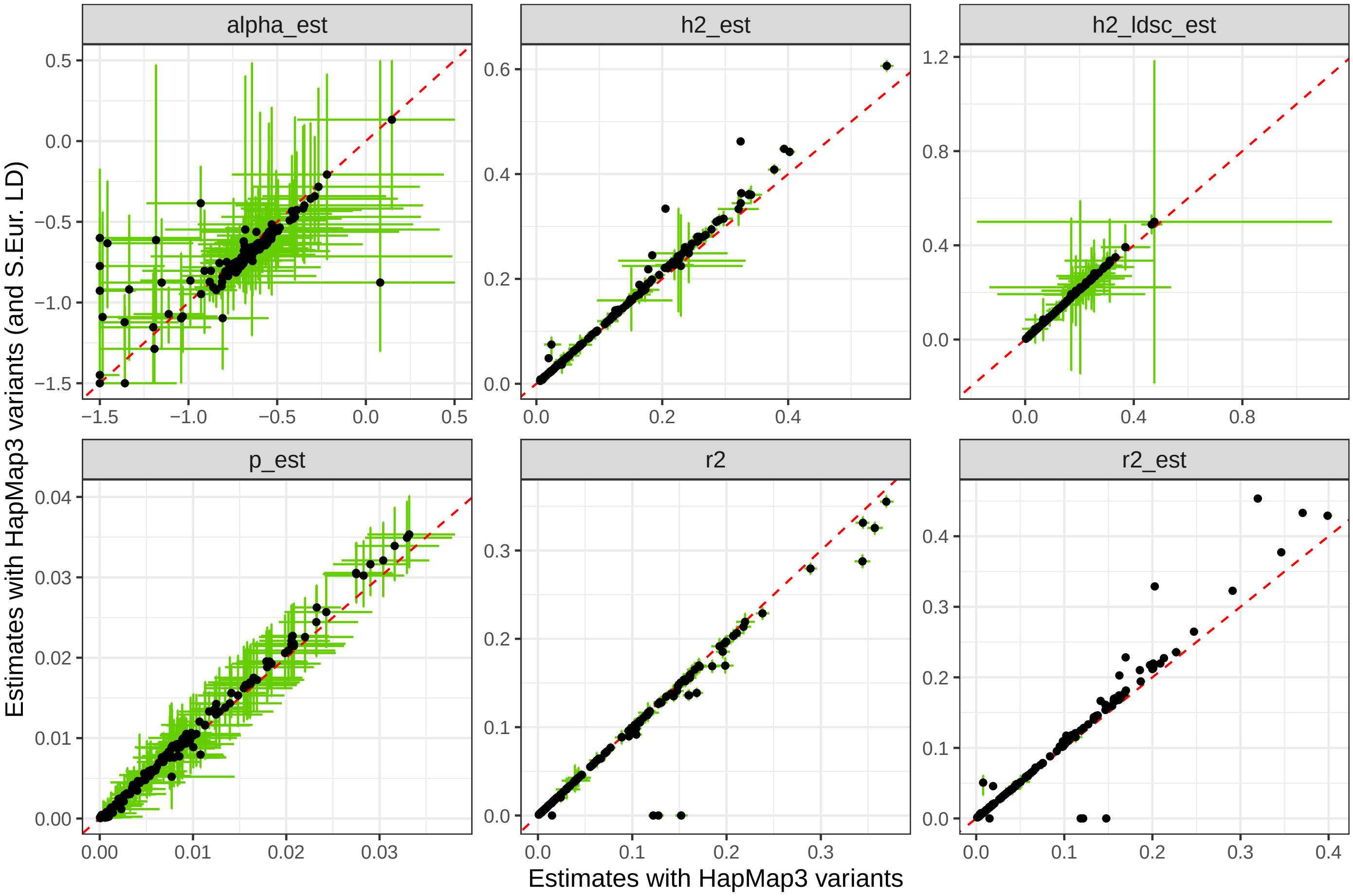


Figure S19: LDpred2-auto estimates for UKBB phenotypes with either a N.W. European or a S. European ("altpop") LD reference. Only 154 phenotypes with more than 25 chains kept when using the N.W. European LD reference are represented here. Red dashed lines represent the 1:1 line. The 95% confidence interval for the LDpred2-auto estimate (in green) is obtained from the 2.5% and 97.5% quantiles of all the estimates from the iterations (after burn-in) of the chains kept. The 95% confidence interval for $r^2$ in the test set is obtained from bootstrap.

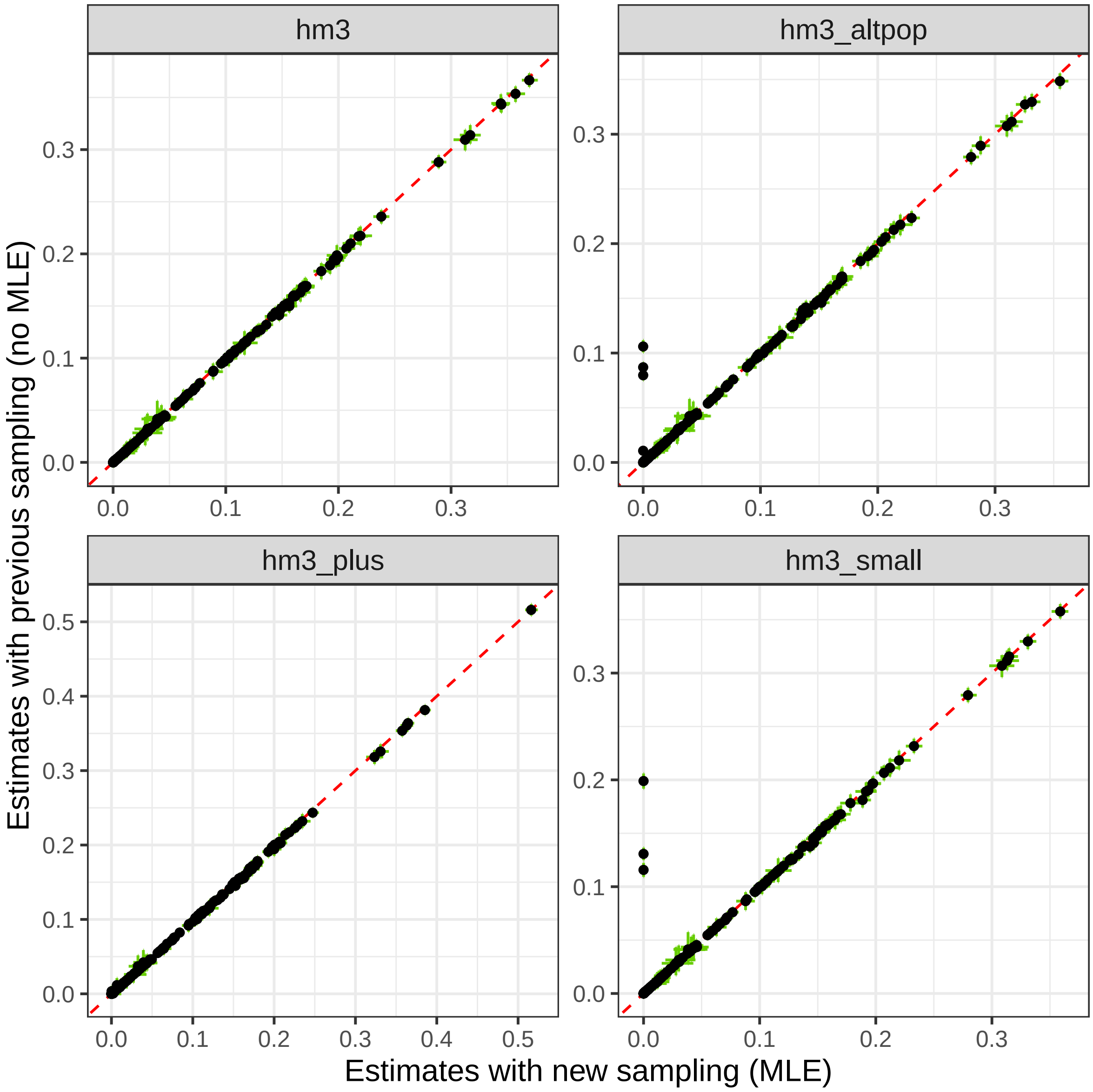


Figure S20: $r^2$ estimates (in the test set) for UKBB phenotypes, using LDpred2-auto with either the new sampling (MLE, 3-parameter model) or the previous one (2-parameter model). Red dashed lines represent the 1:1 line. The 95% confidence interval for $r^2$ in the test set is obtained from bootstrap.

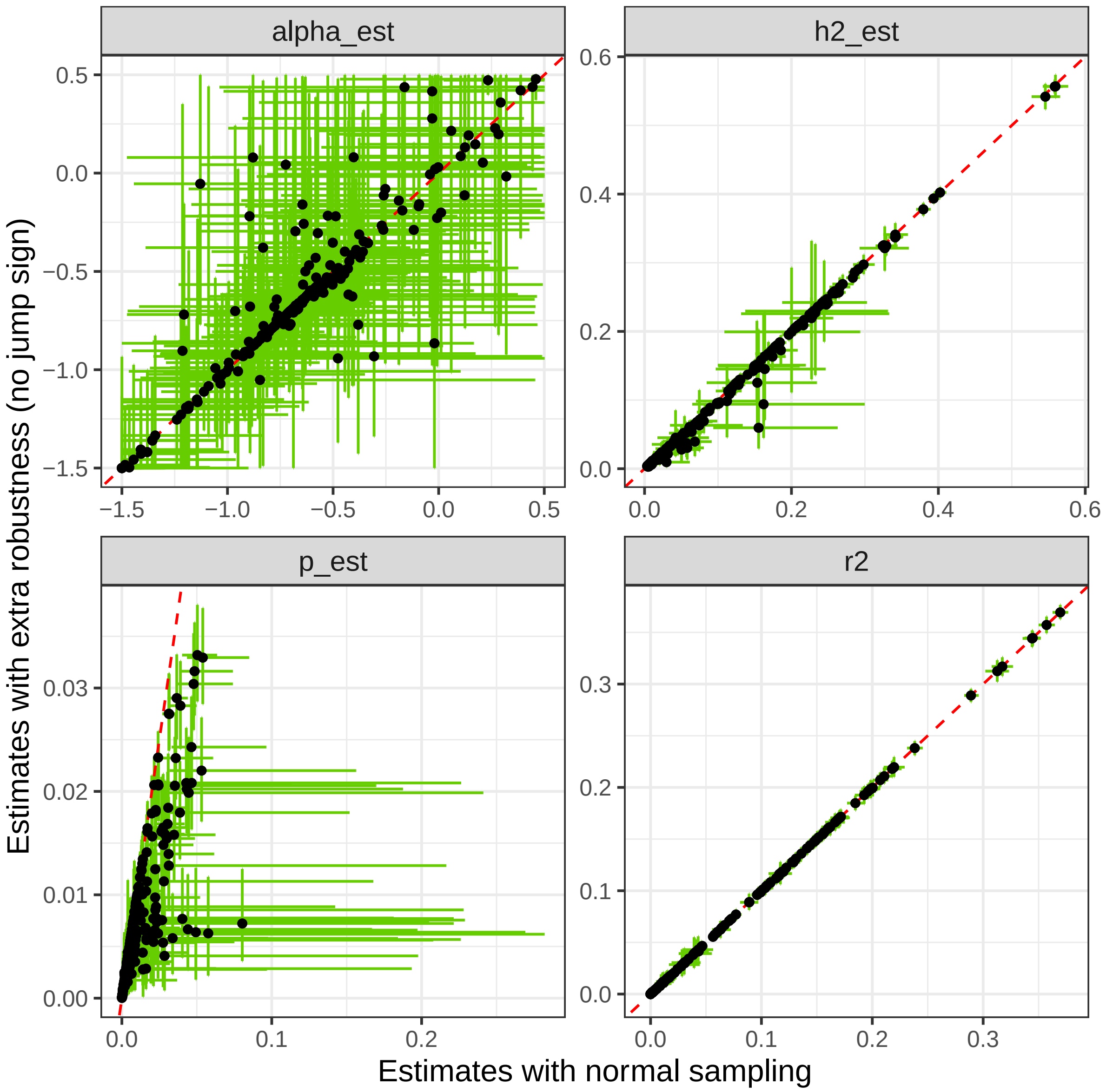


Figure S21: LDpred2-auto estimates for 248 UKBB phenotypes, with or without option ‘allow_jump_sign’ enabled. Red dashed lines represent the 1:1 line. The 95% confidence interval for the LDpred2-auto estimate (in green) is obtained from the 2.5% and 97.5% quantiles of all the estimates from the iterations (after burn-in) of the chains kept. The 95% confidence interval for $r^2$ in the test set is obtained from bootstrap.

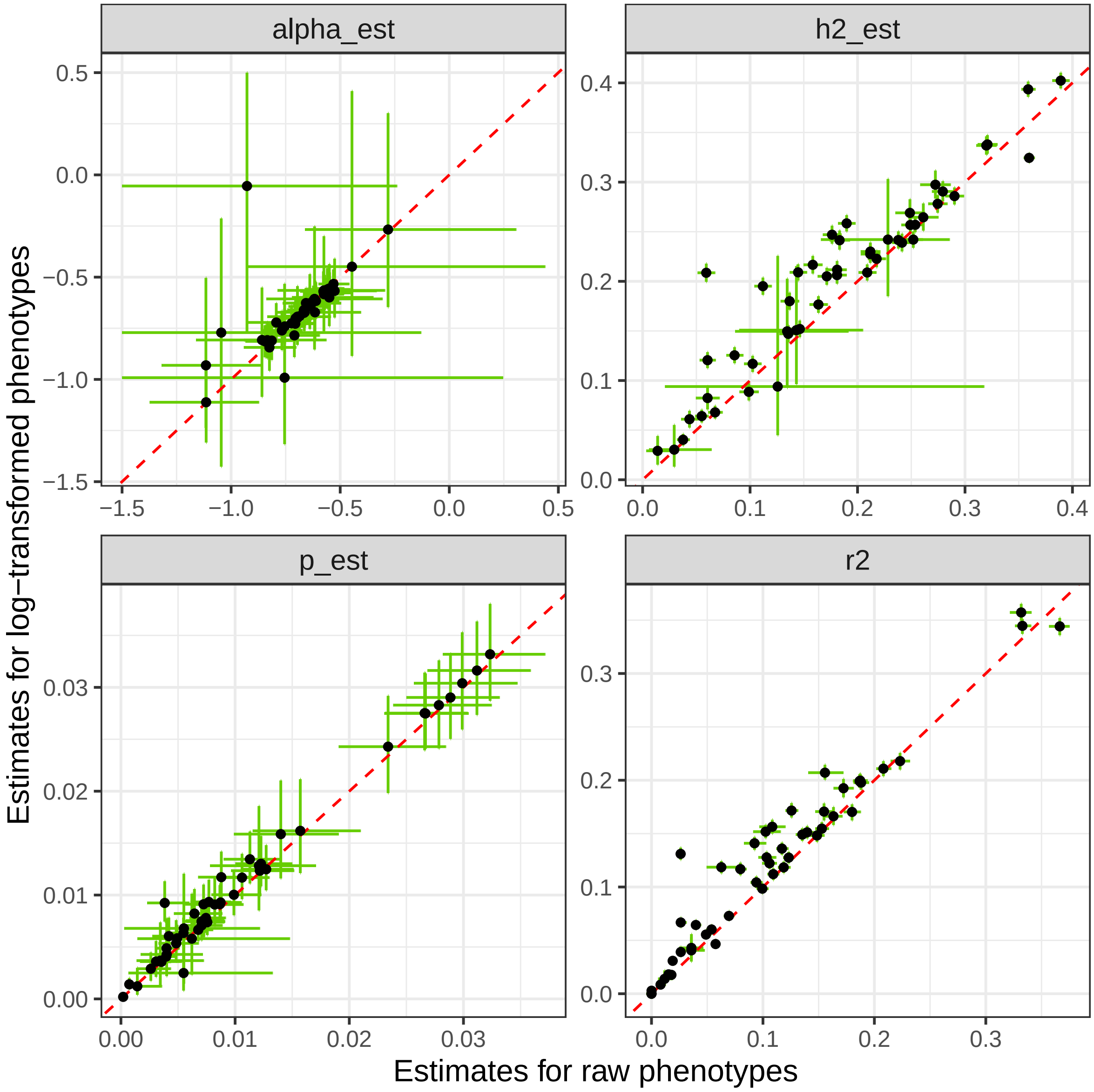


Figure S22: LDpred2-auto estimates for 49 log-transformed UKBB phenotypes, versus for their raw versions. Red dashed lines represent the 1:1 line. The 95% confidence interval for the LDpred2-auto estimate (in green) is obtained from the 2.5% and 97.5% quantiles of all the estimates from the iterations (after burn-in) of the chains kept. The 95% confidence interval for $r^2$ in the test set is obtained from bootstrap.

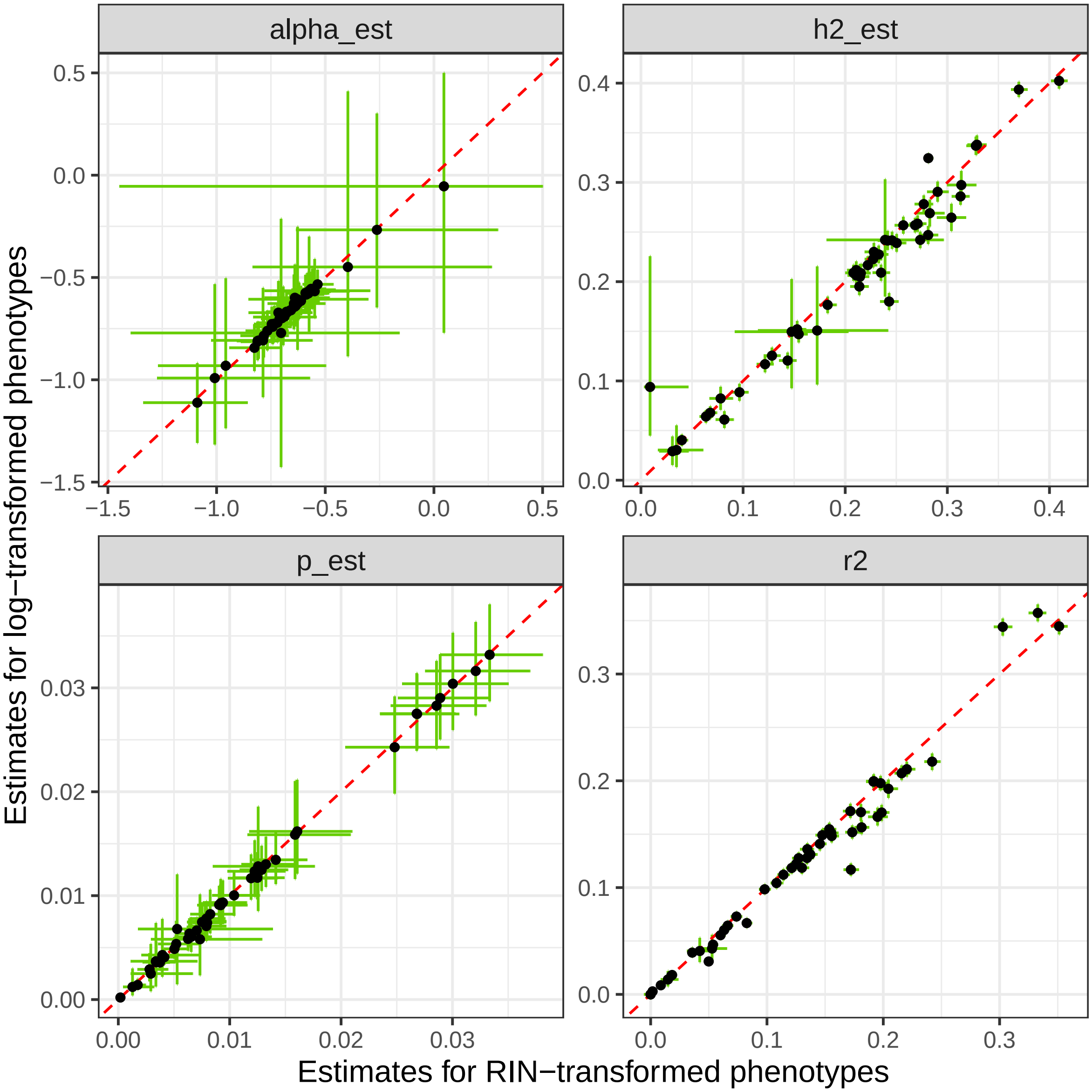


Figure S23: LDpred2-auto estimates for 49 log-transformed UKBB phenotypes, versus for their rank-based inverse normal (RIN) transformed versions. Red dashed lines represent the 1:1 line. The 95% confidence interval for the LDpred2-auto estimate (in green) is obtained from the 2.5% and 97.5% quantiles of all the estimates from the iterations (after burn-in) of the chains kept. The 95% confidence interval for $r^2$ in the test set is obtained from bootstrap.

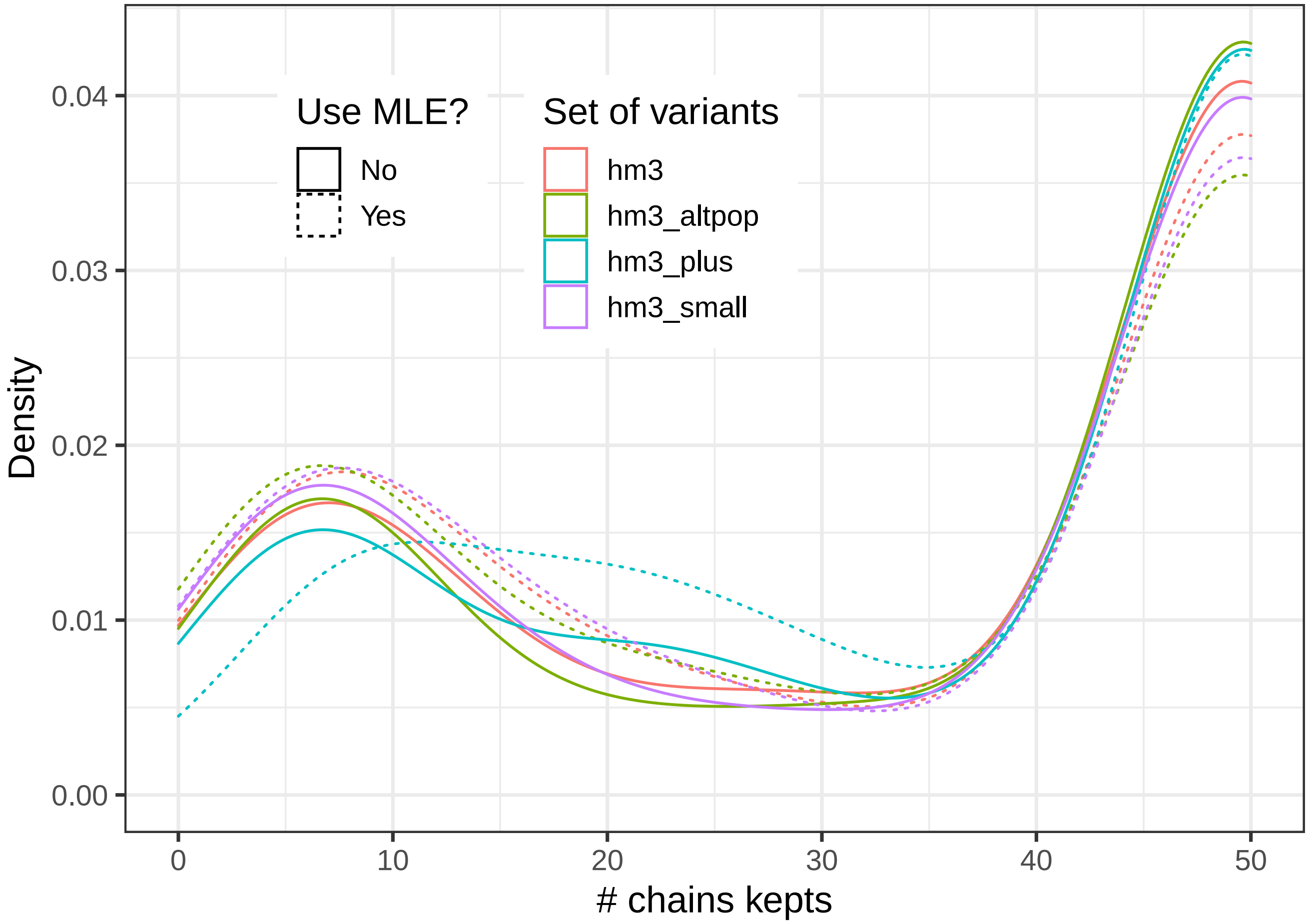


Figure S24: Distribution of the number of LDpred2-auto chains kept across 248 UKBB phenotypes. “Use MLE” corresponds to using the new 3-parameter model and sampling scheme (Methods).

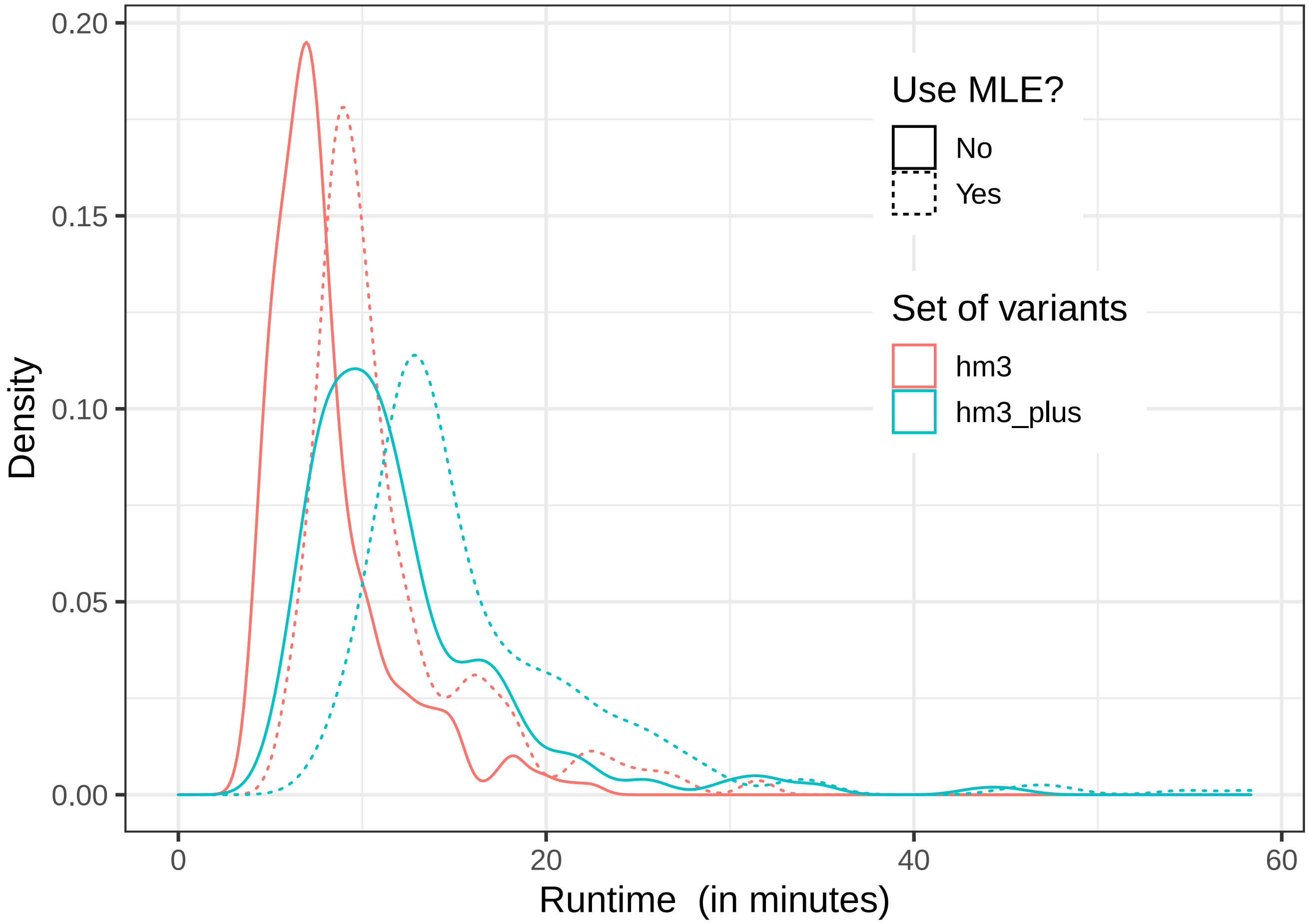


Figure S25: Distribution of LDpred2-auto runtimes across 248 UKBB phenotypes. "Use MLE" corresponds to using the new 3-parameter model and sampling scheme (Methods). For each phenotype, 50 chains are used, parallelized over 13 cores.

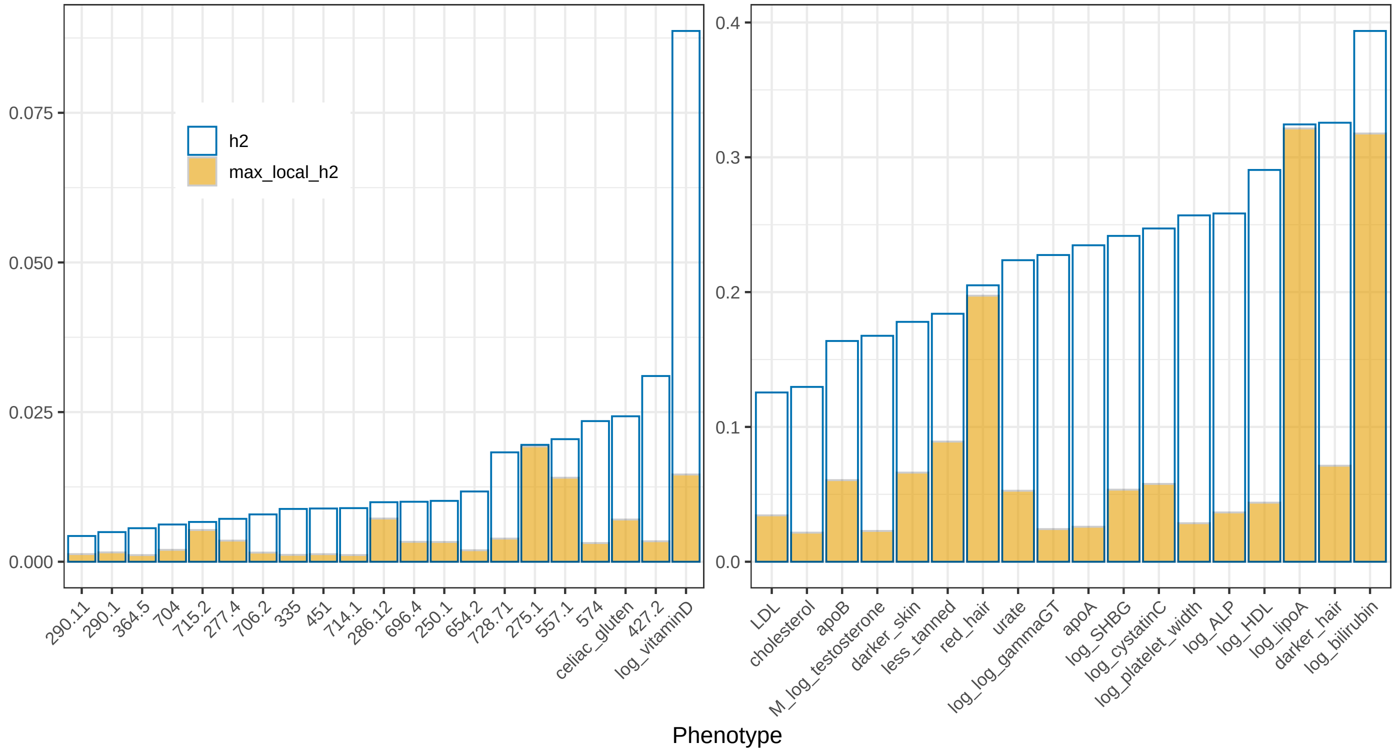


Figure S26: Genome-wide and per-block heritability estimates from LDpred2-auto for UKBB phenotypes. The HapMap3+ variants are used here. The maximum local $h^2$ is the maximum heritability estimate across all 431 independent LD blocks defined for this set of variants. Only phenotypes for which this represents at least 10% of the total heritability are represented.

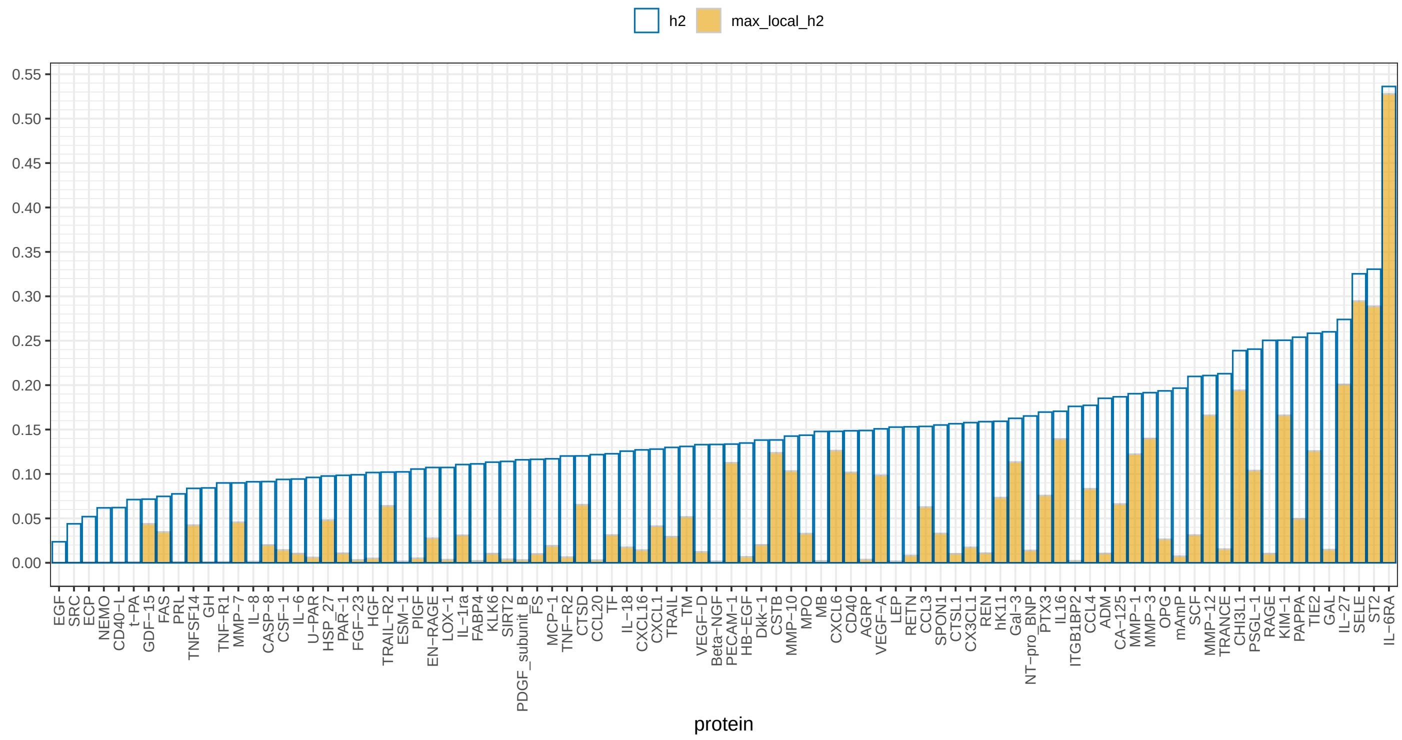


Figure S27: Genome-wide and per-block heritability estimates from LDpred2-auto for 90 protein concentrations[1]. The HapMap3+ variants are used here. The maximum local $h^2$ is the maximum heritability estimate across all 431 independent LD blocks defined for this set of variants.

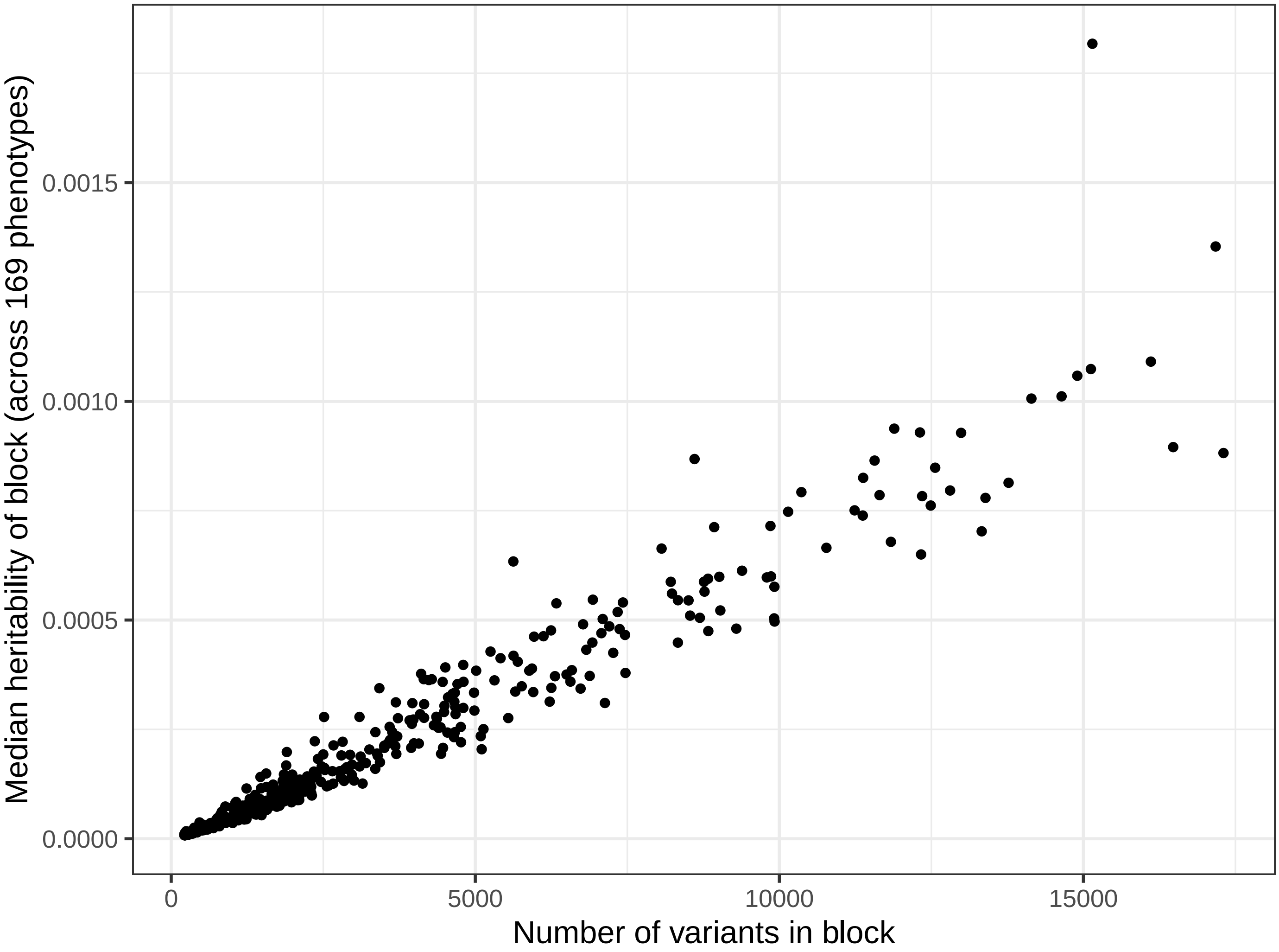


Figure S28: Per-block median heritability across 169 UKBB phenotypes. The HapMap3+ set of variants is used, with 431 independent LD blocks. Only phenotypes with more than 25 chains kept are used here. The top block is on chromosome 6 [22.1-41.4 Mb], which contains the HLA region.

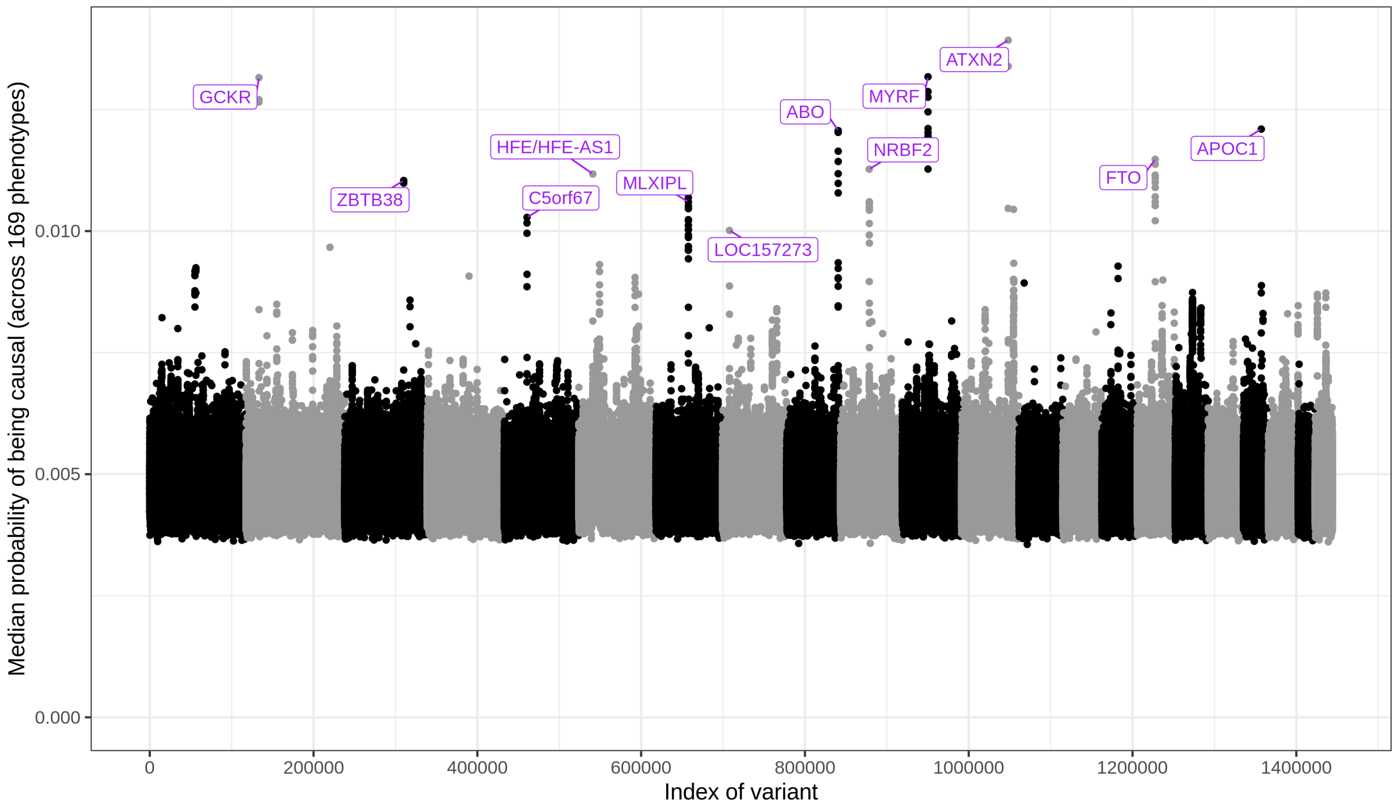


Figure S29: Per-variant median probabilities of being causal across 169 UKBB phenotypes. Variants with a median probability larger than 0.01 were mapped to genes using R package rsnps (only the largest per peak). The HapMap3+ set of variants is used. Only phenotypes with more than 25 chains kept are used here.

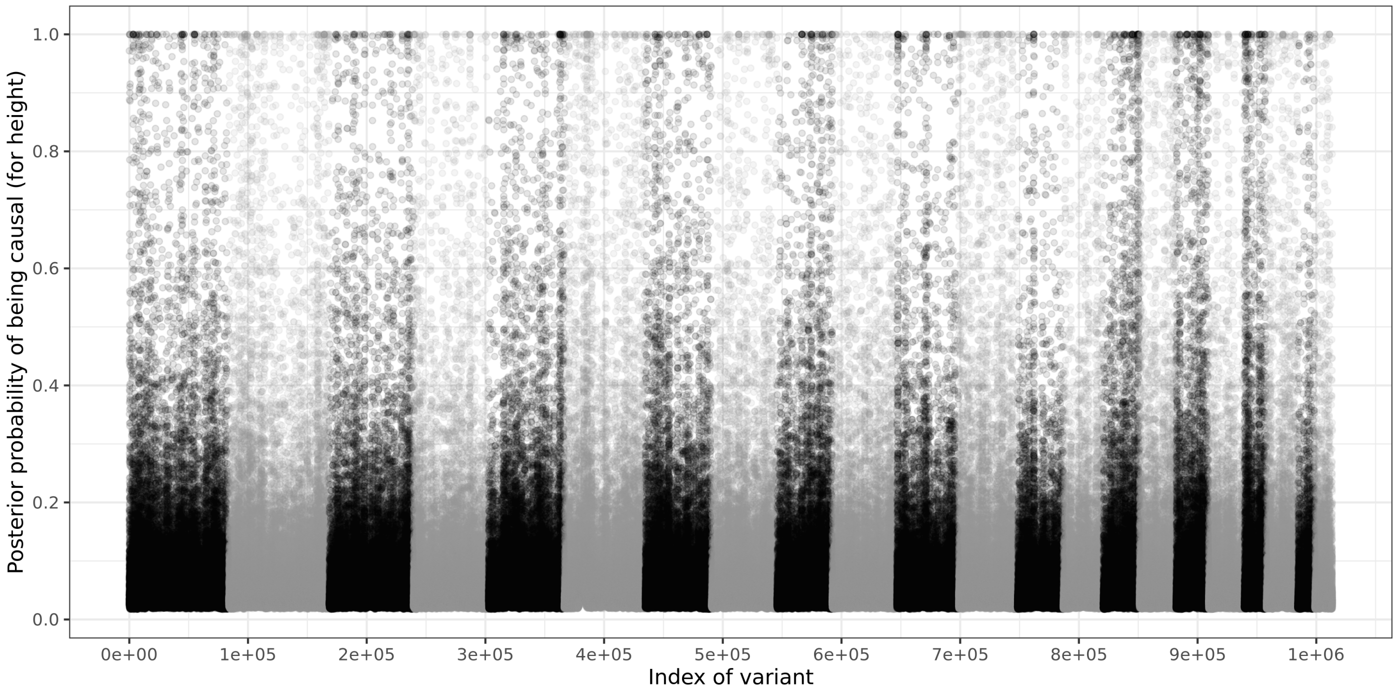


Figure S30: Per-variant probabilities of being causal for height. These are provided by LDpred2-auto when using the external GWAS summary statistics of 1.6M European individuals from Yengo *et al.*[5].

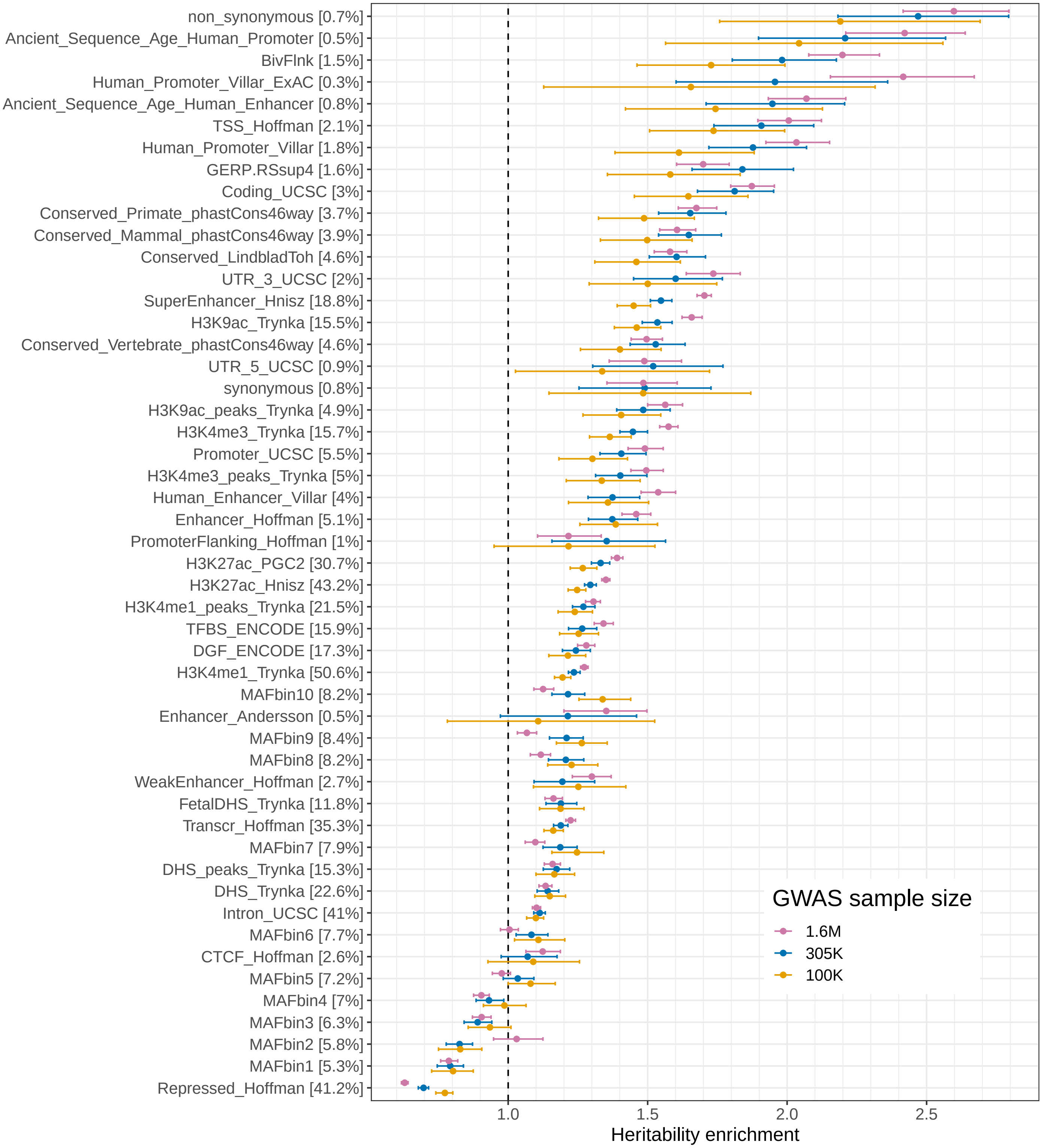


Figure S31: Heritability enrichment from LDpred2-auto for height across 50 functional annotations. The 95% confidence interval for the LDpred2-auto estimate is obtained from the 2.5% and 97.5% quantiles of all the estimates from the iterations (after burn-in) of the chains kept.

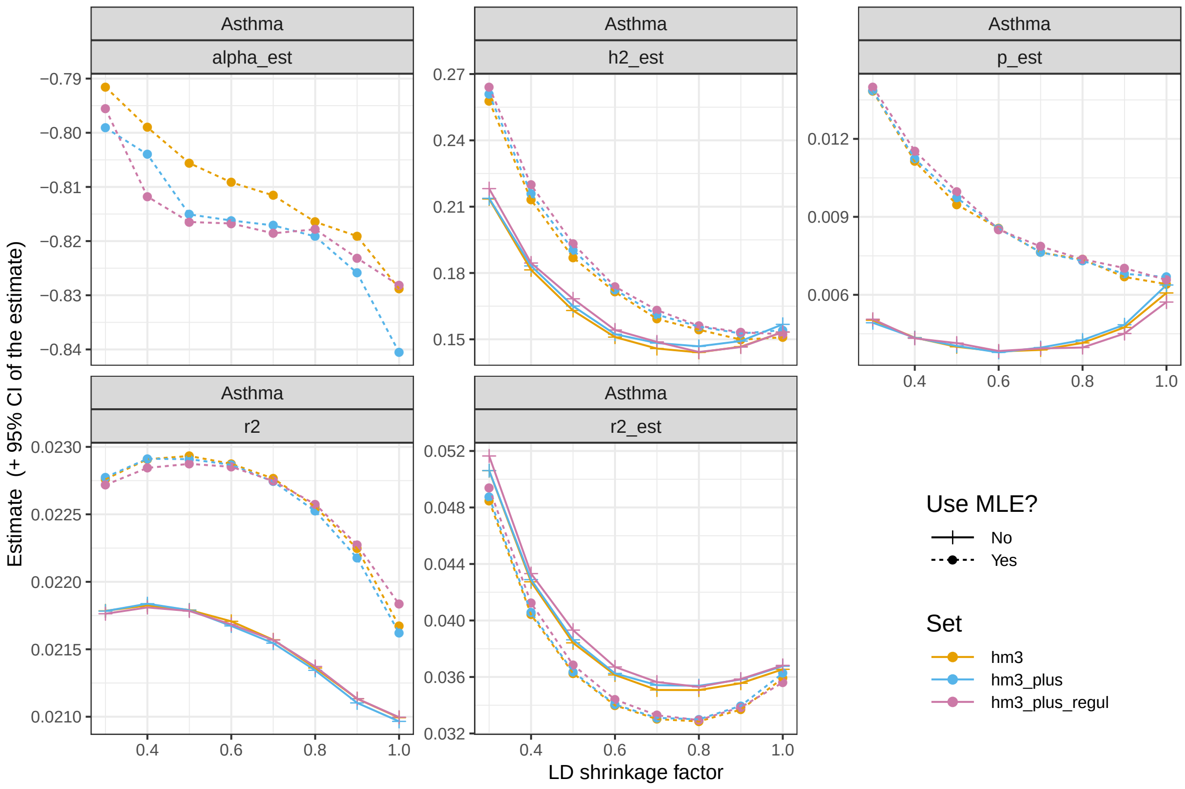


(a) Asthma

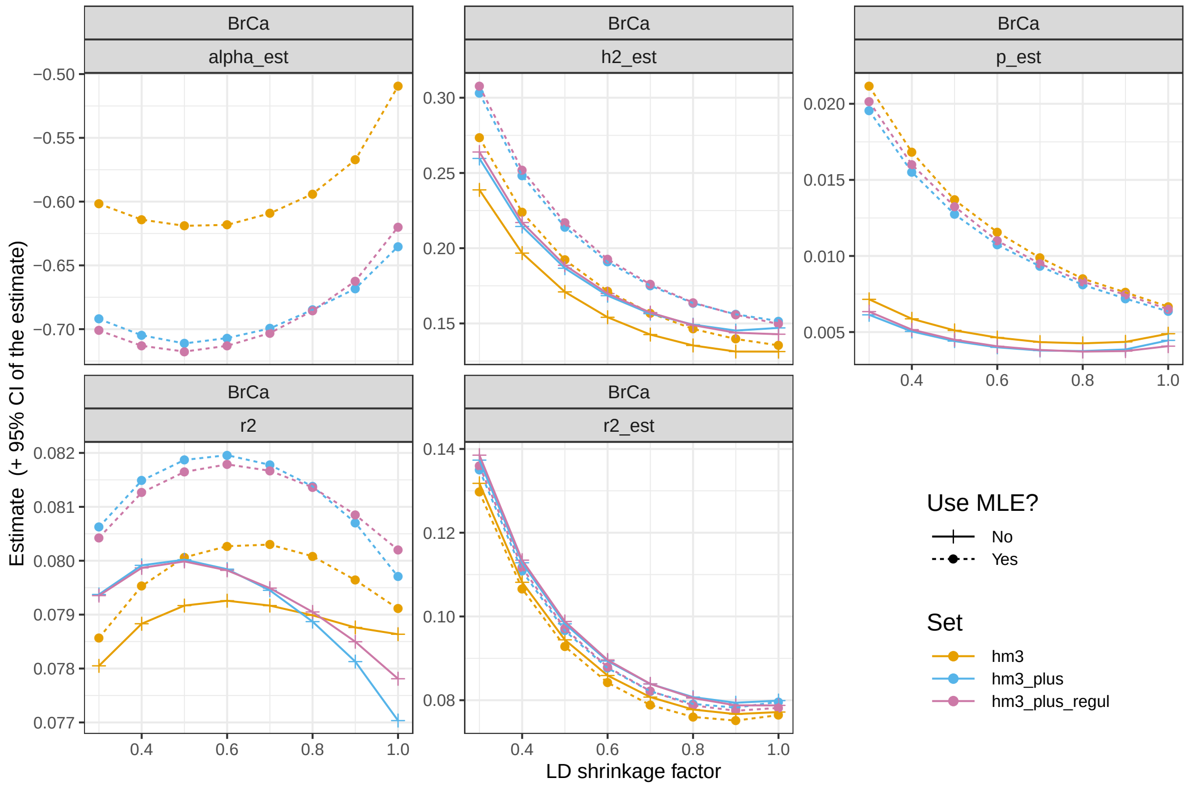


(b) Breast cancer

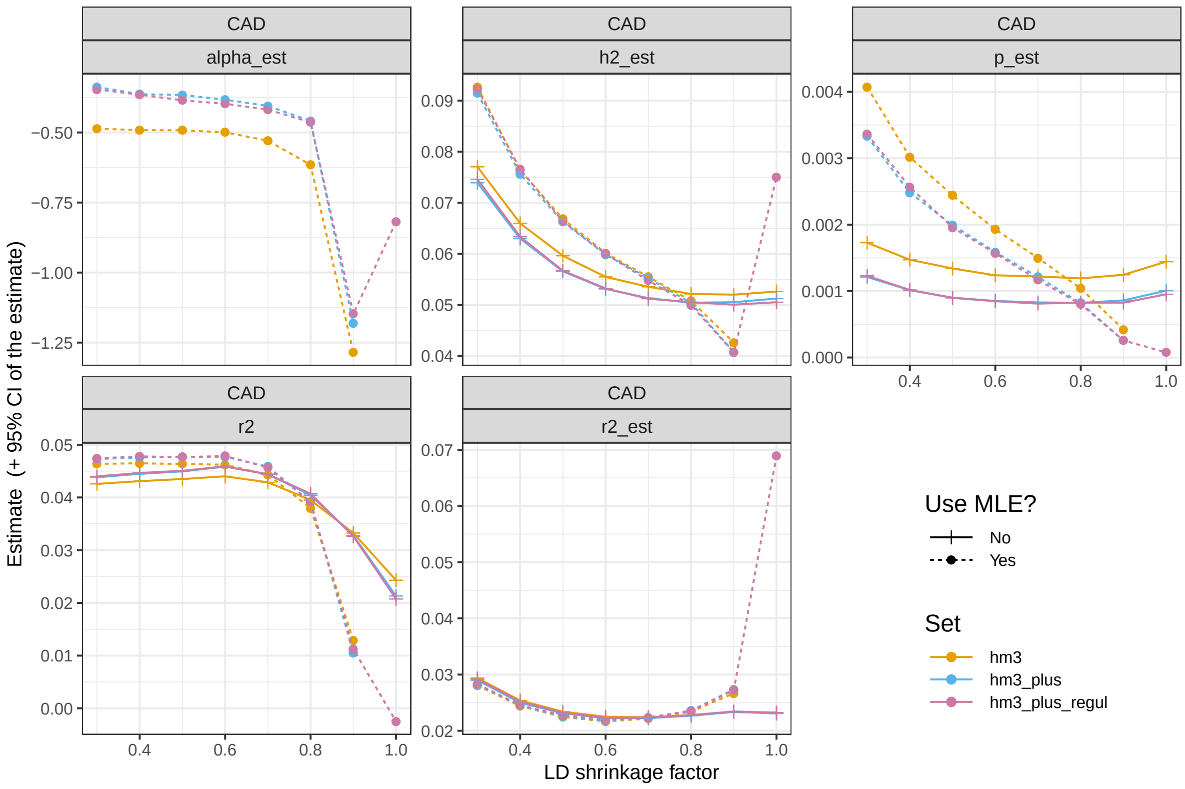


(c) Coronary artery disease

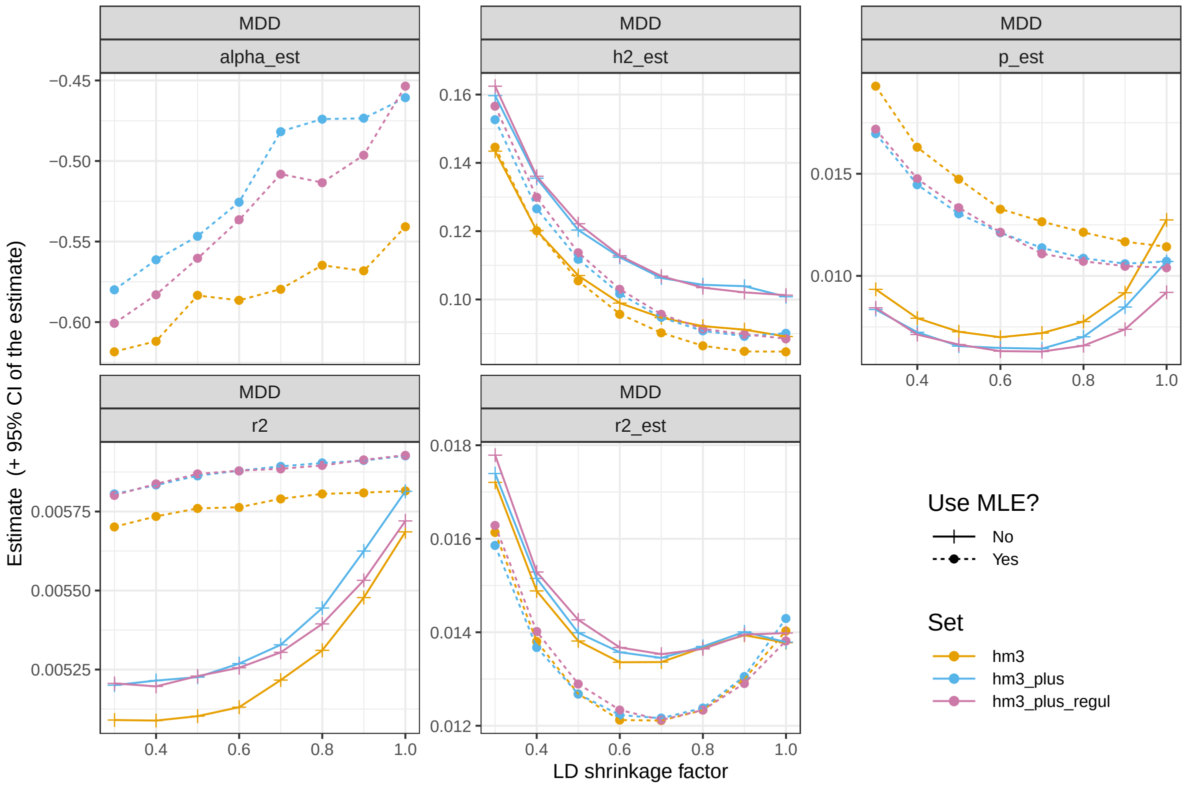


(d) Depression

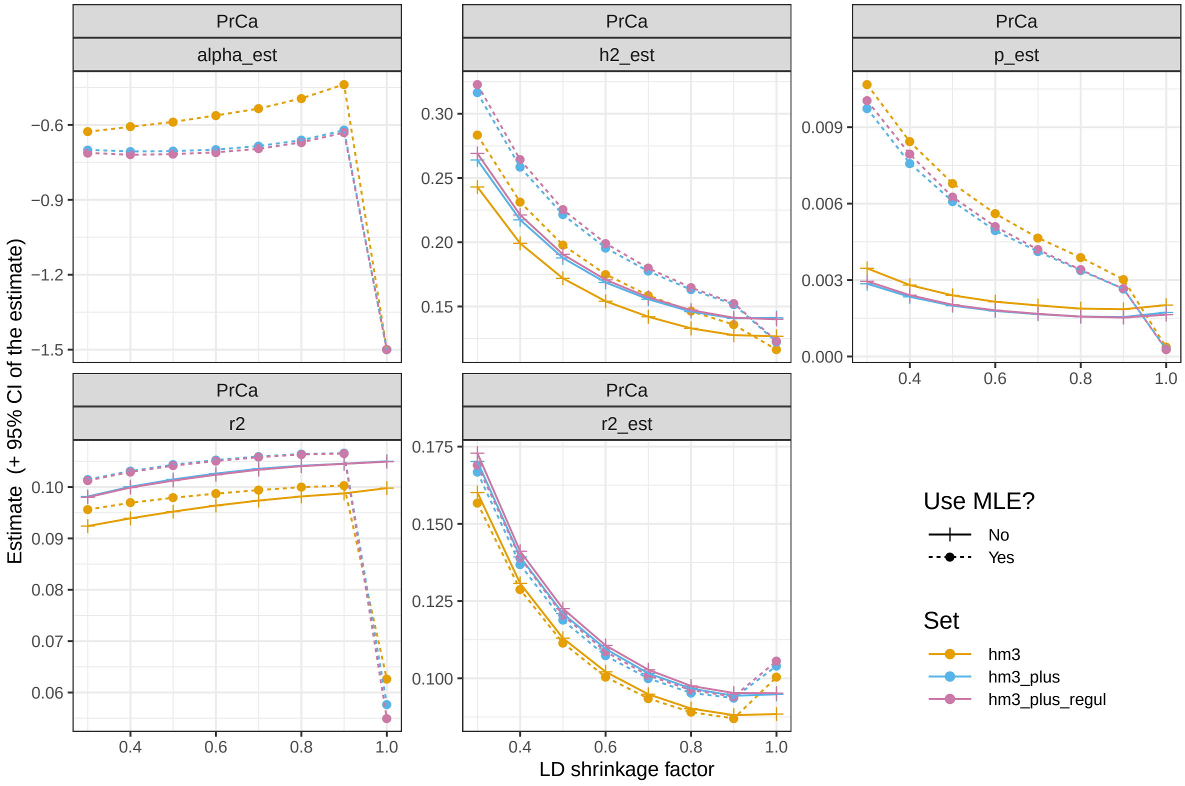


(e) Prostate cancer

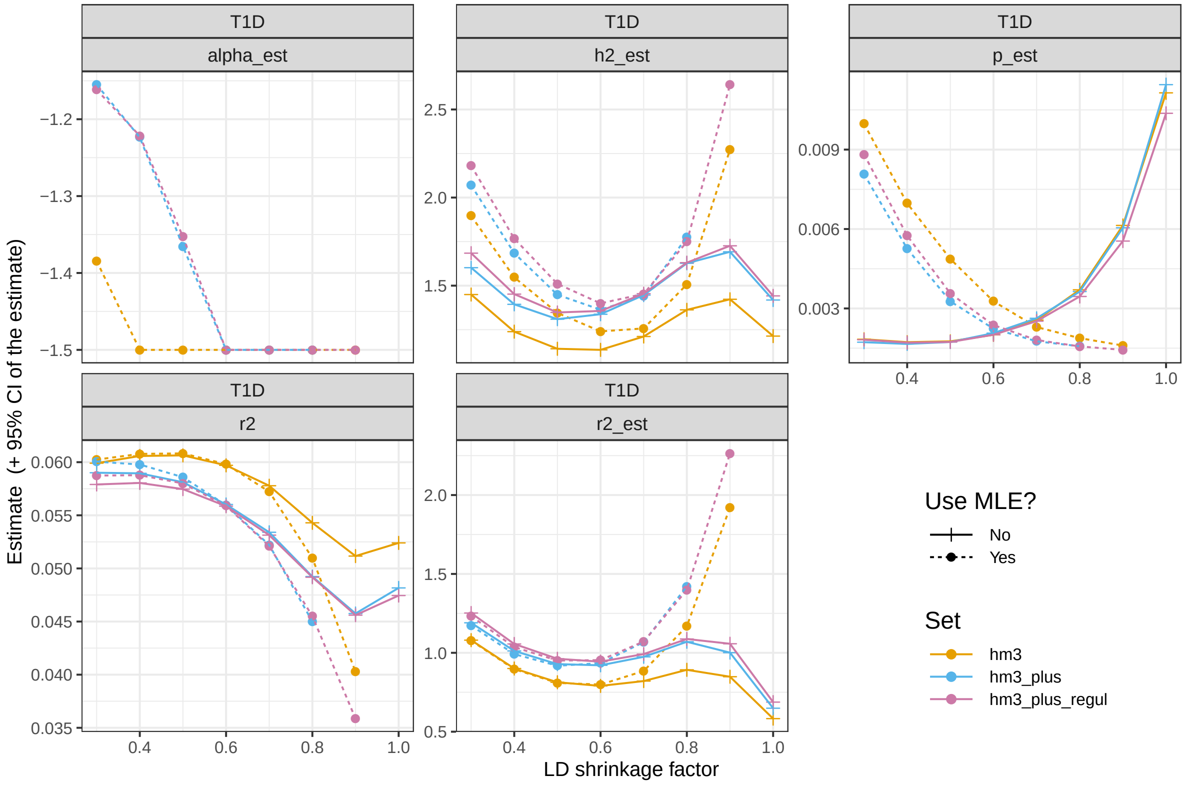


(f) Type 1 diabetes

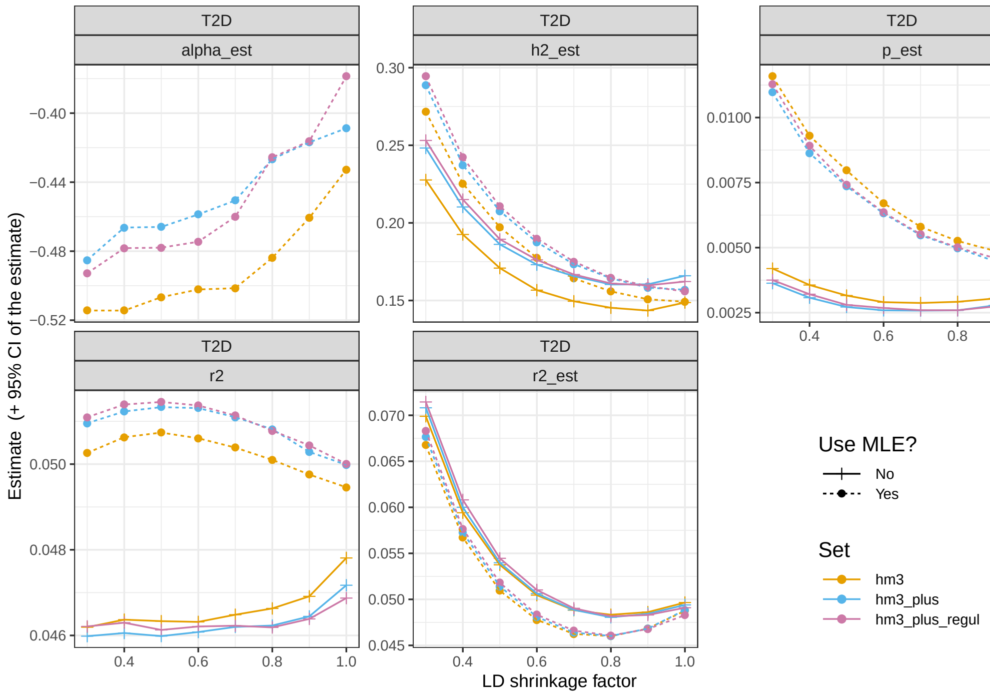


(g) Type 2 diabetes

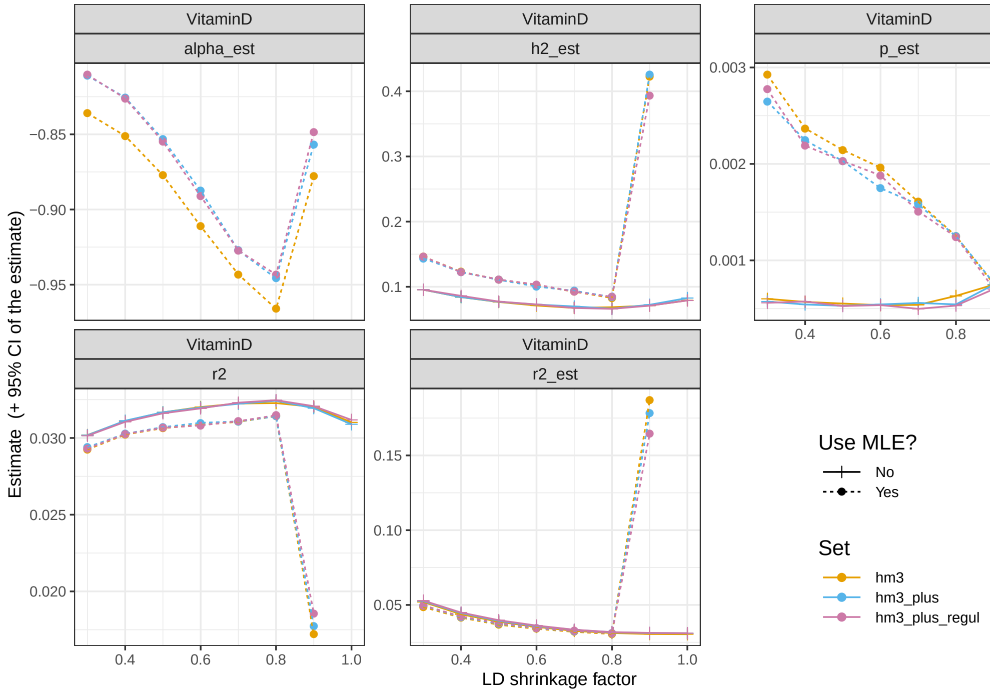


(h) Vitamin D

Figure S32: LDpred2-auto results with external GWAS summary statistics. We run LDpred2-auto using either the HapMap3 or HapMap3+ variants, with either the new or previous sampling and model (via parameter `use_MLE`, where setting to `TRUE` uses the new model), and explore multiple values for parameter `coef_shrink` (multiplicative coefficient for shrinking/regularizing off-diagonal elements of the LD matrix). The UK Biobank is used as test set to compute $r^2$.
